# Rapid depletion of CTCF and cohesin proteins reveals dynamic features of chromosome architecture

**DOI:** 10.1101/2021.08.27.457977

**Authors:** Ning Qing Liu, Mikhail Magnitov, Marijne Schijns, Tom van Schaik, Robin H. van der Weide, Hans Teunissen, Bas van Steensel, Elzo de Wit

## Abstract

The interphase genome is mainly shaped by cohesin-mediated loop extrusion and cohesin-independent compartmentalization. Extrusion is a dynamic process of cohesin loading, loop extension and release. Cohesin release is mediated by WAPL. Loss of WAPL leads to the formation of longer loops and counters compartmentalization. The dynamics of these changes in chromosome organization have been unclear. We have used acute depletion of WAPL to show that within six hours cohesin accumulates at CTCF-bound loop anchors and extended loops are formed. When we deplete WAPL and CTCF simultaneously, new loops are formed between active genes. Surprisingly, active gene clustering is independent of cohesin. Stabilization of cohesin on chromatin leads to a decrease in compartmentalization, which is rapidly restored by depletion of cohesin. Our analyses show that loop extrusion counters compartmentalization and plays a central role in many aspects of chromosome organization.

**HIGHLIGHTS:**

1. Cohesin accumulates at CTCF-mediated chromatin loop anchors following WAPL depletion.
2. Actively transcribed genes form long-range gene clusters independent of the cohesin complex.
3. Plumes are a novel architectural feature of juxtaposed DNA formed by cohesin at open chromatin islands.
4. Chromosome compartmentalization can be uncoupled from nuclear lamina interactions.

## INTRODUCTION

Higher-order genome organization facilitates many fundamental nuclear processes, such as DNA replication, DNA repair and gene expression. Within the nucleus, chromosomes occupy their own specific volumes known as chromosome territories (Cremer and Cremer, 2010). Each individual chromosome is further segregated into active (A) and inactive (B) compartments (Lieberman-Aiden et al., 2009; Simonis et al., 2006; Stevens et al., 2017). Active A compartments are enriched for transcribed genes and active histone modifications such as H3K4me3 and H3K27ac. On the other hand, B compartments tend to be gene-poor and are enriched for histone marks associated with transcriptional repression such as H3K9me3, and are located close to the nuclear periphery (Kind et al., 2015). Compartments are generally subdivided into multiple self-interacting regions called topologically associating domains (TADs) that represent a collection of transiently and stably formed chromatin loops (Dixon et al., 2012; Nora et al., 2012; Rao et al., 2014). In recent years, the mechanisms underlying different levels of genome organization have been extensively studied, uncovering the molecular roles of the key architectural proteins.

The cohesin complex has emerged as one of the major players in interphase genome organization. Cohesin is a ring-shaped protein complex consisting of the subunits SMC1A, SMC3, RAD21 (SCC1), and either SA1 (STAG1) or SA2 (STAG2). This complex is bound to CTCF binding sites genome-wide (Parelho et al., 2008; Wendt et al., 2008) and brings two distal regions in close proximity by forming a chromatin loop. Cohesin and CTCF binding sites are enriched at TAD boundaries to insulate neighboring regions from each other (Dixon et al., 2012). Acute depletion of CTCF or cohesin leads to loss of most TAD boundaries and chromatin loops (Nora et al., 2017; Wutz et al., 2017). The CTCF-mediated chromatin loops are dominantly formed between two convergently oriented CTCF binding sites. The formation of these loops can be disrupted by deleting or inverting one of the paired CTCF motifs (Guo et al., 2015; Sanborn et al., 2015; de Wit et al., 2015), indicating the functional importance of CTCF convergence in loop formation. The specific orientation of CTCF sites that loop together over hundreds of kilobases can be explained by the loop extrusion model (Fudenberg et al., 2016; Sanborn et al., 2015). Recent single-molecule imaging experiments have shown that human cohesin can indeed extrude DNA *in vitro* (Davidson et al., 2019; Kim et al., 2019).

Loop extrusion is considered a highly dynamic process that consists of three steps: (1) loading of cohesin onto chromatin, (2) extrusion of chromatin to form loops, and (3) release of cohesin from chromatin in order to reinitiate the extrusion process. Loading of cohesin is promoted by the cohesin loading complex consisting of NIPBL and MAU2 (also called SCC2 and SCC4). Release of cohesin from chromatin is catalyzed by WAPL. Knock-out of the *Nipbl* or *MAU2* genes leads to a genome-wide loss of chromatin loops and TADs due to a failure to load cohesin (Haarhuis et al., 2017; Schwarzer et al., 2017). Loss of WAPL results in an increased loop size, a decrease of intra-TAD interactions and a loss of compartmentalization (Haarhuis et al., 2017; Wutz et al., 2017). From this it was proposed that loop extrusion counters compartmentalization. Furthermore, the loading and release cycle, collectively referred to as cohesin turn-over, performs a crucial role in transcriptional regulation. Acute depletion of the WAPL protein results in a redistribution of cohesin from binding sites of cell-type specific transcription factors to CTCF binding sites, leading to a decrease in expression for cell-type specific genes (Liu et al., 2021). These results highlight the importance of WAPL-mediated cohesin release in the loop extrusion process.

Recent studies have shown through protein structure analyses, imaging and 3D genome analysis that the N-terminal region is crucial for the role of CTCF in loop formation. Mutating two amino-acids within or deleting the N-terminus of the CTCF protein leads to a genome-wide loss of all CTCF-anchored chromatin loops (Li et al., 2020; Nora et al., 2020). In addition to CTCF, RNA polymerase II is also thought to affect the cohesin positioning at the actively transcribed gene body (Busslinger et al., 2017; Heinz et al., 2018). However, whether and how transcription affects 3D genome organization is a matter of on-going debate. Cells treated with α-amanitin show little effect on the compartment level (Palstra et al., 2008). More recently, Hi-C experiments following depletion of RNA polymerase I, II, and III also revealed only minor effects on 3D genome organization at the TAD level (Jiang et al., 2020).

Rapid inactivation of proteins is essential to study changes in chromosome organization at high temporal resolution and thereby separate direct effects from indirect effects. By acutely depleting WAPL, CTCF and cohesin we have identified cohesin-dependent and -independent mechanisms driving 3D genome organization. We find that in the absence of WAPL and CTCF, new chromatin loops are formed by actively transcribed genes. Depletion of WAPL and CTCF, followed by depletion of cohesin shows that these active gene clusters are not dependent on cohesin. Depletion of WAPL shows a rapid decrease in compartmentalization, which can be reverted by sequential depletion of WAPL and cohesin. These results conclusively show that compartment interactions are counteracted by cohesin-mediated loop extrusion, validating computational models for 3D genome organization (Nuebler et al., 2018). Our experiments show the power of acute protein depletion in the study of chromosome organization, to assess existing models and identify novel features of chromosome architecture.

## RESULTS

### WAPL prevents cohesin accumulation at CTCF-loop anchors

We and others have previously shown that stabilization of cohesin leads to the formation of extended CTCF-anchored chromatin loops. What has been unclear so far is the time frame within which this reorganization occurs. Our previously published WAPL-AID line (Liu et al., 2021), from which WAPL can be depleted within 1 hour after addition of the plant hormone auxin (also known as IAA) offers a unique opportunity to analyze the effect of acute WAPL depletion on chromosome organization. We therefore extended our published *in situ* Hi-C dataset to four time points (untreated, 6h IAA, 24h IAA and 96h IAA). First we asked how quickly loops are extended upon loss of WAPL and observed a clear formation of extended loops as soon as 6 hours following IAA treatment (**Figure 1A**). To quantify loop extension at different time points, we used loops identified in a very deeply sequenced Hi-C dataset generated from the wild-type E14Tg2a cells (Bonev et al., 2017), which we define as ‘primary loops’. We used these primary loops to generate all the pairwise combinations of all primary 5’ anchors with all 3’ anchors up to a specified length, which we refer to as the putative ‘extended loops’ (**Figure 1B**). We quantified the average genome-wide contact frequency for both the primary and extended loops using Aggregate Peak Analysis (APA), and observed a gradual decrease in contact frequency for primary loops following WAPL depletion (**Figure 1C**). The APA confirmed that extended loops were almost fully formed after formed 6 hours after starting the IAA treatment and remained up until the last measurement at 96h (**Figure 1A and 1C**). Another striking feature of WAPL loss is the decrease in intra-TAD interactions. To determine the dynamics of intra-TAD interactions we performed an Aggregate TAD Analysis (ATA) using high-resolution TADs previously identified in wild-type mESCs (Bonev et al., 2017). This also revealed that intra-TAD interactions diminish rapidly, while corner peaks remain present (**Figure 1C**). These results show that loss of WAPL leads to a rapid and lasting reorganization of the 3D genome.

**Figure 1:**
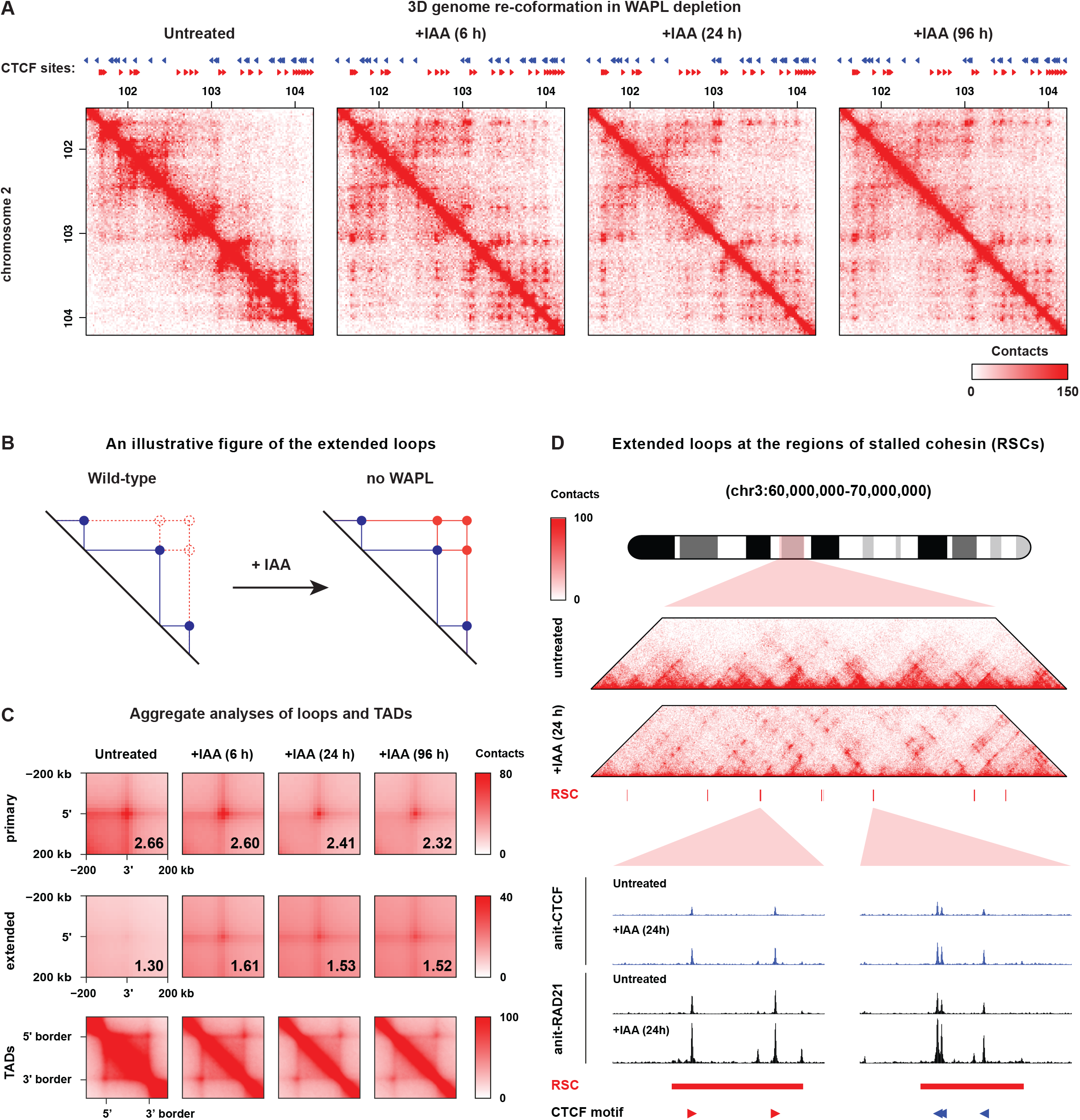
Acute WAPL depletion leads to a rapid reorganization of interphase chromosomes. (A) Hi-C matrix plot at 20kb resolution showing an example locus at chromosome 2. CTCF sites are from CTCF ChIPseq in untreated WAPL-AID cells. (B) An illustrative figure describing the primary and extended loops. (C) Aggregate analyses reveal the genome-wide changes of chromatin loops and TADs upon WAPL depletion. The primary loops and TADs are obtained from Bonev et al. (2017), and the extended loops are predicted from the primary loops base the method described in Haarhuis et al. (2017), with a maximum loop length of 3Mb. (D) An example showing formation of the extended loops between two regions of stalled cohesin (RSCs) reported in Liu et al. (2021). Zoomed in regions of ChIPseq tracks show changes in CTCF and cohesin binding in untreated and WAPL depleted cells. Forward and reverse CTCF binding motifs within the RSC are indicated by red and blue triangles, respectively.

We have previously shown that cohesin is rapidly redistributed upon WAPL depletion. In mESCs we identified 2,789 regions that showed increased cohesin binding, which we termed Regions of Stalled Cohesin (RSCs) (Liu et al., 2021). Inspection of our Hi-C maps suggested that loop anchors often overlap with RSCs (**Figure 1D**). Indeed, of the 12,807 unique loop anchor sites forming the 9,376 primary loops (Bonev et al., 2017), 1,822 overlap with RSCs. Conversely, 58% of RSCs (1,619 out of 2,789) overlap with loop anchors. We tested whether the presence of an RSC at loop anchors differentially affected the response to WAPL depletion. For primary loops, the RSC overlapping loops remain prominent, while the loops that have no RSC present in their anchors become clearly weaker (**Figure S1A, left panel**). Furthermore, the presence of RSCs at the anchors correlates with emergence of extended loops. When both predicted anchors overlap with an RSC, the extended loops are strongly induced upon WAPL depletion. In contrast, loop anchors that do not overlap with RSCs form only very weak extended loops (**Figure S1A, right panel**). These results show that cohesin accumulates at loop anchors, and indicates that the amount of cohesin accumulation determines the looping propensity between two distal loop anchors.

Based on these results we hypothesized that we could predict chromatin interactions purely based on the linear position of the RSCs. To test this, we generated a set of putative loops in silico based on the pairwise combination of RSC locations (**Figure 2A**). APA analysis revealed a clear increase in contact frequency following WAPL depletion (**Figure S1B**), indicating that RSCs are indeed sites where extended loops are formed. Next, we wondered whether the newly formed loops adhered to the CTCF convergency rule. To investigate this, we stratified the RSCs based on unique CTCF motif orientation. Note that we excluded RSCs that contain CTCF-binding sites with a non-unique orientation from this analysis. We observed that the predicted convergent loops interact most strongly upon WAPL depletion (**Figure 2B**, top row). However, pairwise combinations between tandemly oriented RSCs show an average contact frequency that is almost as high as that of the convergently oriented pairs (see quantification in **Figure 2B**). These results clearly indicate that loss of WAPL rapidly induces the formation of loops between CTCF/cohesin sites that do not necessarily adhere to the convergency rule.

**Figure 2:**
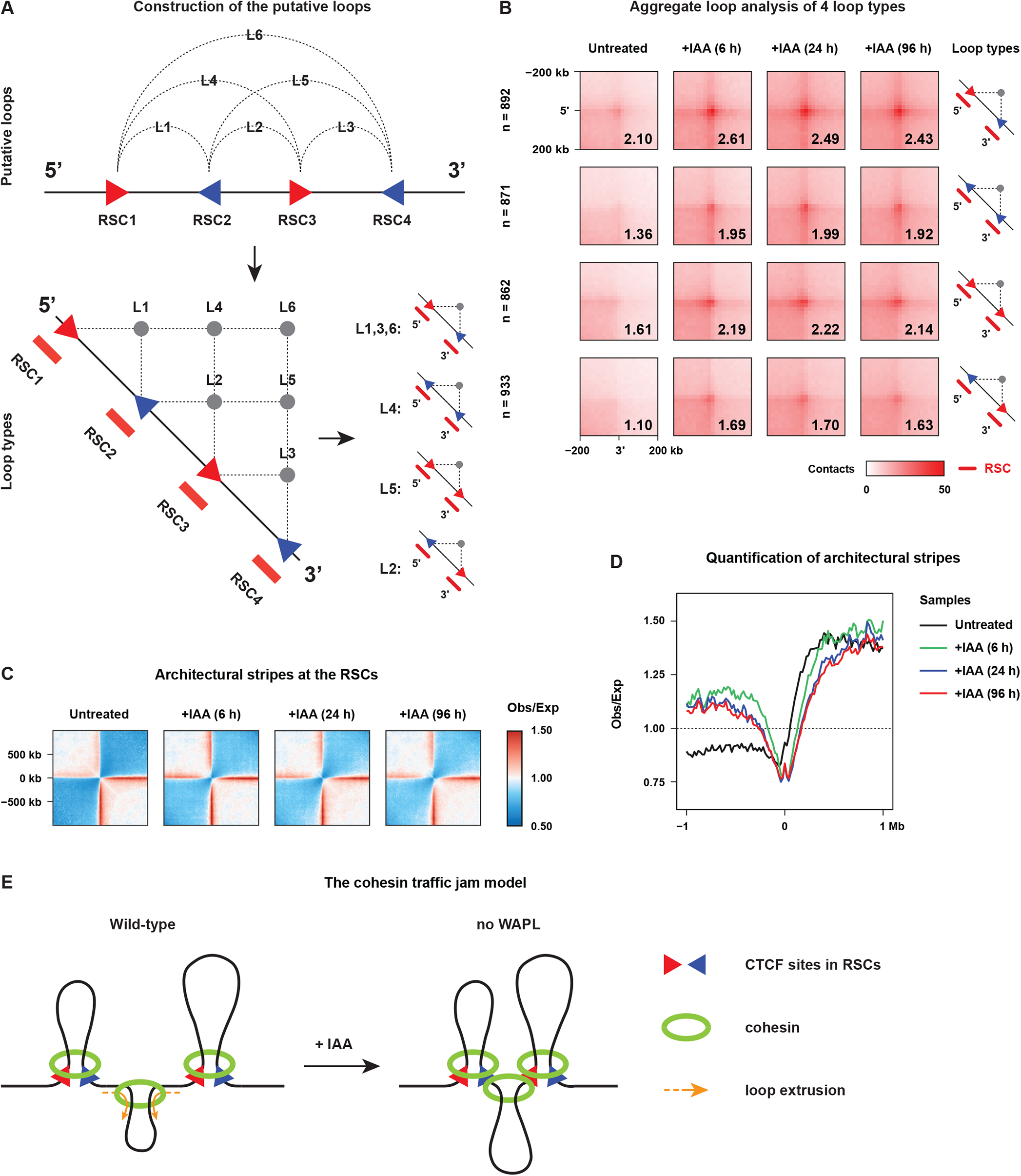
Genomic regions where cohesin accumulates are anchors of chromatin loops. (A) An illustrative figure describing construction of the putative loops from RSCs. (B) Aggregate loop analysis quantifies intensity of the chromatin loops between two RSCs containing CTCF sites with unique orientations. The distance of two RSCs is restricted up to 6Mb. (C,D) Aggregate region analysis measures observed over expected interactions of architectural strips anchored by RSCs. (E) A model explains the formation of loops between non-convergent CTCF sites and counter-stripes.

CTCF-anchored chromatin loops are thought to be formed through cohesin-mediated loop extrusion. One principle feature in Hi-C maps that is indicative of loop extrusion are so-called stripes (Barrington et al., 2019; Vian et al., 2018). Stripes can be visualized by aligning Hi-C data on features that act as stripe anchors, such as CTCF binding sites (Hansen et al., 2019). Therefore, we aligned our Hi-C data on RSCs that have a unique orientation. We found that in untreated cells a clear stripe can be seen emanating from the position of the RSC (**Figure 2C,D**). Interestingly, when WAPL is depleted a stripe in the opposite orientation, or counter-stripe, is formed. This observation is consistent with a model in which accumulation of cohesin at convergent CTCF sites serves as a roadblock for the cohesin extruded in the opposite direction, which may also explain the formation of loops involving two non-convergent CTCF sites (**Figure 2E**).

### Actively transcribed genes form long-range loops in the absence of WAPL and CTCF

In the absence of WAPL, cohesin rapidly accumulates at CTCF sites (Liu et al., 2021) and loop anchors. Busslinger et al. showed that in the absence of both WAPL and CTCF cohesin accumulates at active genes in so-called ‘cohesin islands’ and is this relocation is driven by transcription (Busslinger et al., 2017). However, whether these cohesin complexes are involved in chromatin looping is not known. To study this we generated a double degron cell line from which both WAPL and CTCF could be acutely depleted. To this end we fused an AID-mCherry to the C-terminus of the endogenous CTCF protein in the WAPL-AID-GFP line (**Figure S2A**). We refer to this double-degron line as the WAPL/CTCF-AID line. WAPL and CTCF are completely depleted within 6 hours after the addition of IAA to the culture medium (**Figure S2B**). We performed calibrated ChIP-seq in these cells for WAPL, CTCF, and RAD21 in untreated cells and following 24 hours of IAA treatment. The acute depletion led to a clear genome-wide loss of WAPL and CTCF binding in these cells (**Figure S2C and S2D**). We also observed a clear diminishment of RAD21 occupancy at CTCF binding sites as a direct consequence of global CTCF loss (**Figure S2D**). Importantly, alignment of RAD21 ChIP-seq data at TSS peaks on H3K4me3 peaks from untreated WAPL-AID cells, showed that RAD21 binding to active TSSs is largely unaffected upon the loss of WAPL and CTCF (**Figure S2E**). We confirmed the accumulation of cohesin in cohesin islands in mESCs following WAPL and CTCF depletion (**Figure S2F**). These results show that depletion of WAPL and CTCF is complete and consistent with previously published findings.

To determine what the effect of WAPL/CTCF depletion is on chromosome organization we generated Hi-C maps in untreated, 6h, 24h and 96h treated samples. Genome-wide quantification of relative contact probability (RCP) shows a gradual increase in long-range contacts over time following WAPL/CTCF depletion, in contrast to WAPL only depletion where no further increase is observed after 24 hours of IAA treatment (**Figure S3A**). As expected, we found that loops and TADs rapidly disappear upon WAPL and CTCF depletion (**Figure 3A**). The APA and ATA analyses showed that these architectural features are lost in a genome-wide fashion (**Figure S3B**), in correspondence with observations in CTCF depletion experiments (Nora et al., 2017; Wutz et al., 2017).

**Figure 3:**
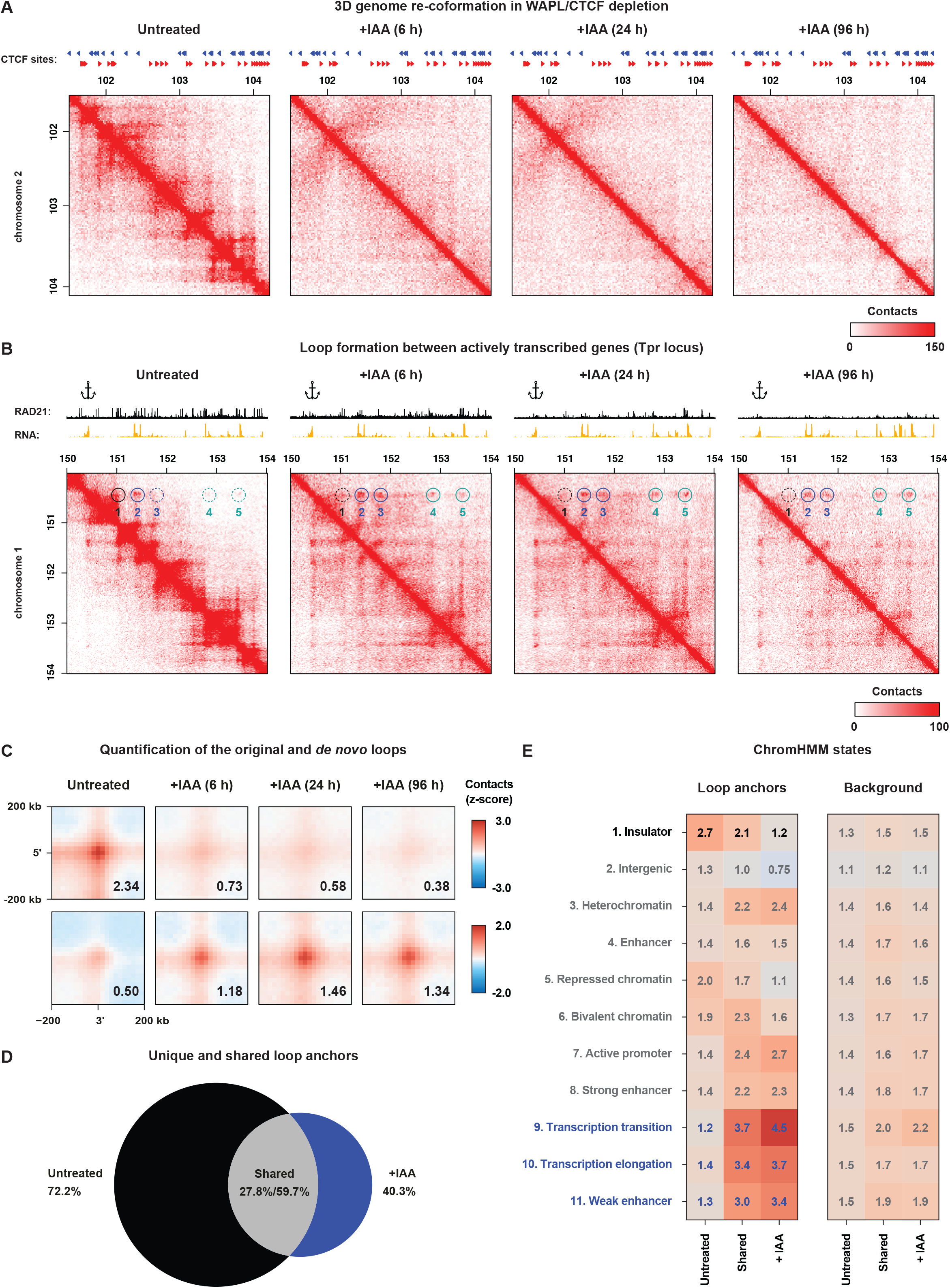
Acute double depletion of WAPL and CTCF results in loop formation between highly transcribed genes. (A) Hi-C matrix plot of WAPL/CTCF depletion experiment; same region and resolution as shown in Figure 1A. CTCF sites are from CTCF ChIPseq in untreated WAPL-AID cells. (B) Hi-C matrixplot of the *Tpr* locus at 20kb resolution showing that the chromatin loops are formed between highly expressed genes after WAPL/CTCF depletion. From the *Tpr* gene (anchor), five loops are identified in the four time points. The solid circles indicate that the loops are detected in the specified time point, while dashed circles mark the loops identified only at the other time points. (C) Aggregate loop analysis quantifies the contact frequency in untreated cells (0h, original loops) and IAA treated cells (6, 24, and 96h, *de novo* loops) using the ChromoSight loop caller, respectively. (D) Venn diagram showing the unique and shared loop anchors in the untreated and IAA treated cells. (E) ChromHMM state enrichment of the three different types of loop anchors indicated in (D).

Although CTCF-anchored loops and TADs were diminished, new chromatin loops were formed in the absence of WAPL and CTCF (**Figure 3B, Circle 3-5**). To characterize these, we performed *de novo* loop calling using ChromoSight, a more sensitive loop caller in high-resolution Hi-C datasets with a comparable precision as HiCCUPS (Matthey-Doret et al., 2020). We find a decrease in the number of loops from 4,804 in the untreated cells to 2,716 in the treated cells for the WAPL/CTCF-AID cells (**Figure S3C**). In contrast, WAPL only depletion leads to an increase in the number of loops from 4,416 in the untreated control to 14,167 in the treated cells (**Figure S3C**). To quantify loop strength for the loops called in the untreated and the treated conditions we performed an APA (**Figure 3C**) and find a clear decrease for the loops identified in untreated cells and an increase for the loops in cells without WAPL and CTCF. This result indicates that, there is a genome-wide shift from one loop type to a novel loop type. To determine what chromatin features loop together in the absence of WAPL and CTCF we functionally annotated these loop types. To this end we first stratified the loop anchors into three classes: loop anchors found i) only in the untreated conditions (**untreated**), ii) in both treated and untreated condition (**shared**) and iii) found only in the treated condition (**+IAA**) (**Figure 3D**). We intersected the loop anchors for these three classes with ChromHMM data generated for mouse ESCs (Pintacuda et al., 2017). Unsurprisingly, the loop anchors specific for the untreated cells strongly overlap with the Insulator class, which is marked by the binding of CTCF. Loop anchors specific for the WAPL/CTCF-depleted cells are not enriched for the Insulator class but are strongly enriched for classes associated with active transcription (**Figure 3E**). These results show that WAPL and CTCF prevent the ectopic clustering of active genes in the 3D genome.

Next, we sought to determine whether this clustering of active genes is the consequence of cohesin accumulation at active genes (Busslinger et al., 2017). To answer this question, we would like to induce active gene clustering, followed by inactivation of the cohesin complex. To achieve this we aimed to independently degrade WAPL/CTCF and the cohesin subunit RAD21. We took advantage of another acute depletion strategy called ‘degradation tag’ (dTAG) (Nabet et al., 2018). In the WAPL/CTCF-AID cell line, we fused an FKBP^F36V^ peptide at the C-terminus of the cohesin subunit RAD21 using our previously published CRISPR-Cas9 strategy (Liu et al., 2021) (**Figure 4A**). Using this “triple degron” cell line, we are able to independently deplete WAPL/CTCF (with IAA) and RAD21 (with dTAG-13) (**Figure 4A,B)**. Western blot analyses confirmed the validity of the two-step depletion strategy (**Figure S4A**). With this two-step degradation system, we can accurately control the timing and order of WAPL/CTCF depletion and cohesin depletion. To dissect formation and maintenance of chromatin loops we performed Hi-C experiments in this triple degron line (**Figure 4B**). As expected, depletion of WAPL/CTCF recapitulated what we had found previously: an increase in contact frequency between loci that are between 1M and 10Mb apart (**Figure S4B**) and a loss of loops and TADs (**Figure S4C**). Depletion of RAD21 has previously been shown to lead to a loss of CTCF-anchored chromatin loops and TADs (JDP et al., 2020; Rao et al., 2017; Wutz et al., 2017), which is recapitulated in our triple degron line (**Figure S4C**). These results show that the triple degron line recapitulates the observations of WAPL/CTCF depletion and previously published RAD21 depletion. To determine whether the loops formed between active genes were dependent on cohesin we performed sequential depletion of WAPL/CTCF and RAD21. By depleting WAPL/CTCF for 24 hours we induced the clustering of active genes, followed by (co-)depletion of RAD21 for three hours. To our surprise, in the absence of cohesin the interactions between active genes are largely maintained (**Figure 4C**), showing that active gene clustering is not entirely dependent on cohesin.

**Figure 4:**
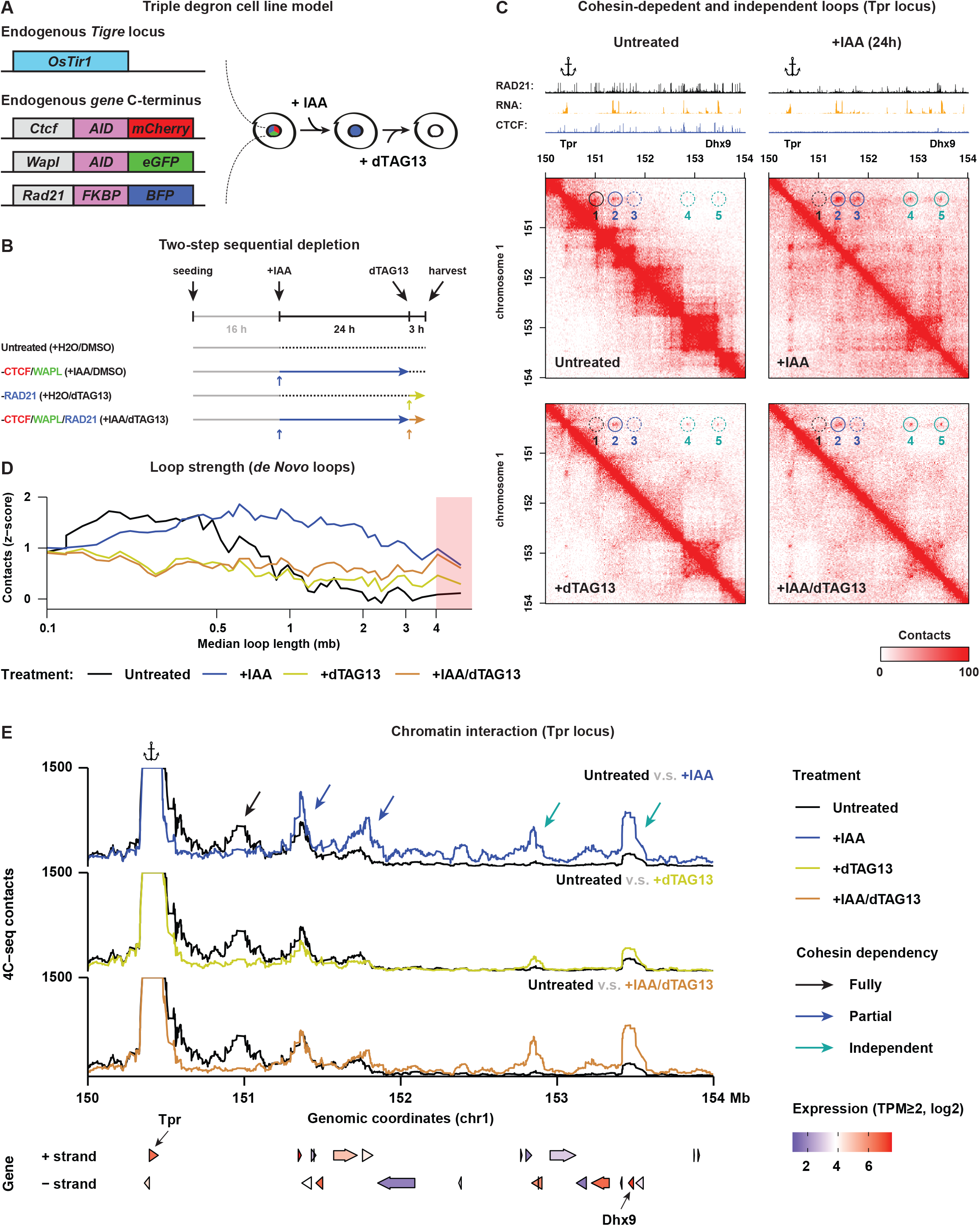
Functional dissection of active gene loops by sequentially depleting WAPL/CTCF and RAD21 using a dual-degradation triple degron cell line. (A) In the OsTir1 E14 mESCs, the endogenous *Wapl* and *Ctcf* gene were tagged with AID–eGFP and AID-mCherry at the C-terminus of WAPL and CTCF, respectively. In this cell line, the endogenous *Rad21* gene was further fused with a FKBP12^F36V^-BFP sequence at the C-terminus of RAD21. In this cell line, we can sequentially deplete WAPL/CTCF and RAD21 by supplementing IAA to the culture medium followed by dTAG13 treatment. (B) A schematic figure describes the strategy of sequential depletion in our experiments. (C) Three types of chromatin loops are observed at the *Tpr* locus. Loop 1 is sensitive to WAPL/CTCF or RAD21 depletion, indicating that it is a CTCF and cohesin dependent loop. Loop 2 and 3 were strengthened or induced through WAPL/CTCF depletion, and reduced by further depletion of RAD21, indicating these two loops are CTCF independent but partially cohesin dependent. Loop 4 and 5 were induced by WAPL/CTCF depletion, which were not reduced after sequentially depleting RAD21, indicating these loops are cohesin independent. (D) Quantification of the *de novo* loops in the triple depletion experiment. The loops are divided into 50 equally sized bins based on their length and the average z-score normalized contact frequency is plotted. (E) Validation of *de novo* loop formation at the *Tpr* locus shown in (C) using 4C-seq.

If the loops between active genes are independent of cohesin, we may expect to observe them also in the absence of cohesin. Indeed, we can observe loops between active genes in an example locus following depletion of RAD21 only (**Figure 4C**). Note that these are the same loops that are induced by WAPL/CTCF depletion. However, the interaction between *Tpr* and *Dhx9* is stronger in the cells in which WAPL and CTCF are pre-depleted, compared to cells in which only RAD21 is depleted (**Figure 4C**). We sought to determine whether stronger interactions between de novo formed loops in WAPL/CTCF pre-depleted cells compared to non-pre-depleted cells following RAD21 depletion is systematic. We performed APA on distance normalized contact matrices, to correct for the substantially different contact probabilities between the four Hi-C datasets (see **Methods**). Quantification of the APAs showed that the longest *de novo* loops (> 3Mb) indeed were stronger in the absence of cohesin following WAPL/CTCF pre-depletion (**Figure 4D**, **Figure S4D**). As a control we performed APA on the original loops and found that the RAD21 only depleted loops responded similar to the two-step WAPL/CTCF+RAD21 depletion (**Figure S4E**).

To confirm the maintenance of active gene looping following cohesin depletion we used 4C-seq (Splinter et al., 2012) to perform a targeted conformation capture approach. We designed viewpoint primers on the *Tpr* promoter and used HindIII digestion and in-nucleus ligation. Our 4C data confirmed that active gene loops were maintained in the absence of cohesin following pre-depletion of WAPL and CTCF (**Figure 4E**). Notably, the longest two loops are weakly induced by single depletion of RAD21 for 3 h, suggesting these long-range gene loops are repressed by cohesin (**Figure 4E**). In summary, our results show that we can separate two mechanisms in the formation of between active genes. Stabilization of cohesin and depletion of CTCF results in the ectopic clustering of active genes (“formation”). Subsequent loss of cohesin, however, does not necessarily lead to a decrease in interactions (“maintenance”). Importantly, although cohesin accumulates at active genes and active genes cluster in the 3D genome, cohesin is only required for creating the spatial proximity in which these loops can form. The actual formation of loops between active genes likely depends on an alternative mechanism.

### Transient architectural features form at small open chromatin islands

Upon inspection of the Hi-C maps for WAPL/CTCF, we observed that at certain sites the signal in the Hi-C maps accumulates perpendicular to the diagonal (**Figure 5A**), but only for the 6h and 24h timepoints. Because the signal seems to emanate from the diagonal we refer to these patterns as “plumes”. When we overlaid regions that encompass plumes with ATAC-seq and H3K4me1 ChIP-seq profiles, we found that plumes are formed at active chromatin domains flanked by large heterochromatic regions (**Figure 5B**). To systematically determine whether regions with a high density of active marks flanked by larger regions nearly devoid of such marks show plume formation, we first identified such regions. For this we used ATAC-seq data to identify regions of high ATAC signal (0.5-1Mb), flanked on either side by a region of low ATAC signal (min. 1Mb). We identified 105 such regions, which we refer to as “open chromatin islands” (OCIs). Next, we aligned the Hi-C signal on these OCIs using Aggregate Region Analysis (ARA). We observed a clear plume pattern following depletion of WAPL/CTCF (**Figure 5C**). Note that when we deplete WAPL alone, we do not observe a plume pattern (**Figure S5A and S5B**). As expected, we also observed a clear enrichment of RAD21 binding at the OCIs, which is drastically decreased after depletion of WAPL and CTCF (**Figure 5B and S5C, left panel**). When WAPL is depleted on its own, CTCF likely restrains the cohesin complex within the range of the OCIs, exemplified by the fact that if we calculate the density of RAD21 signal we find that upon depletion of WAPL this is not affected. However, when we deplete both WAPL and CTCF we see a clear decrease in the signal of RAD21 (**Figure S5C, right panel**).

**Figure 5:**
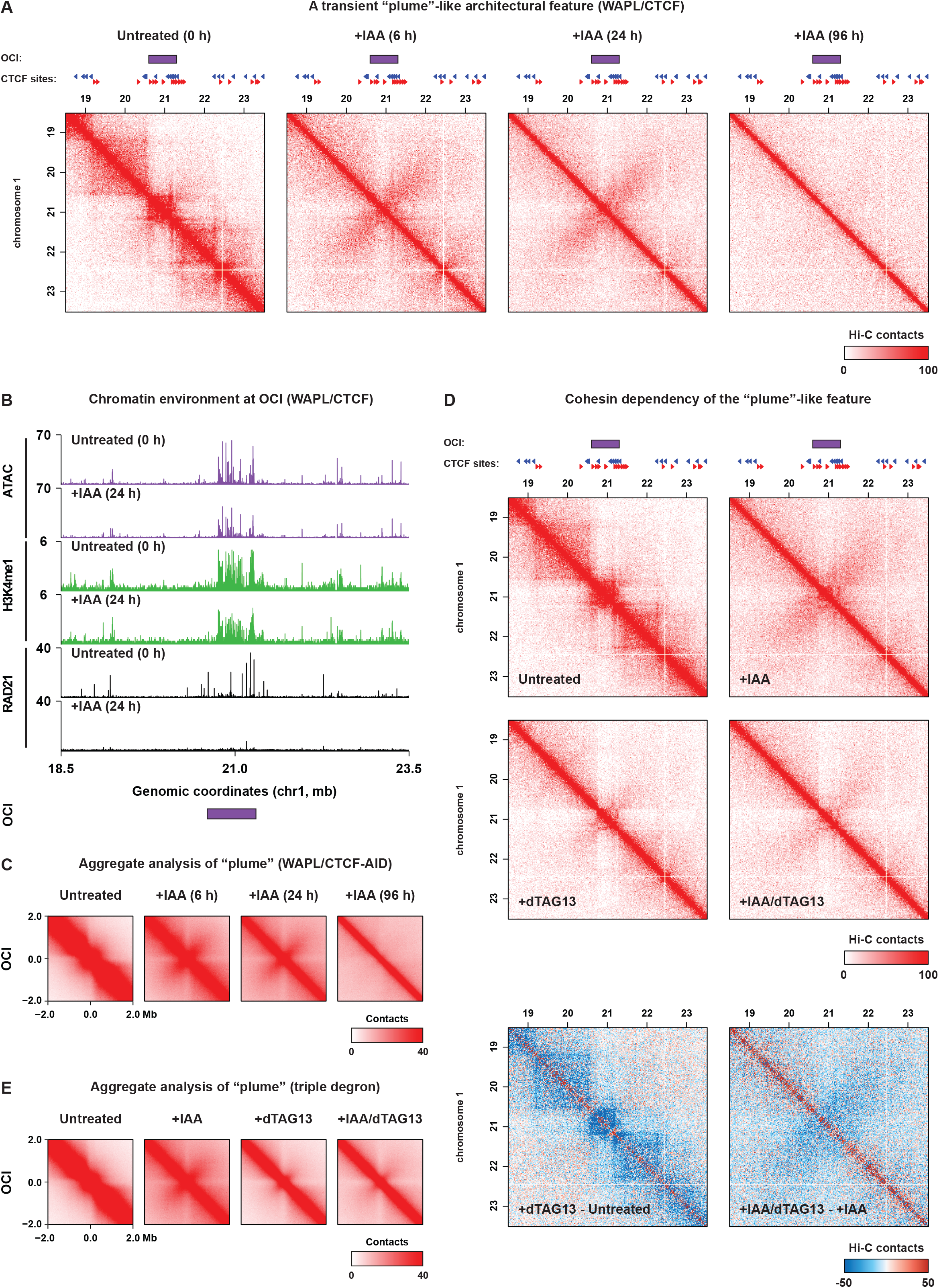
Acute depletion of WAPL and CTCF facilitates transient formation of the “plume”-like architectural features at open chromatin islands (OCIs). (A) Hi-C matrix plot showing a plume example region on chromosome 1 at 20kb resolution. (B) Chromatin environment of an OCI showing a clear enrichment of open chromatin sites, H3K4me1, and cohesin compared to the surrounding regions. After WAPL/CTCF depletion, chromatin accessibility and H3K4me1 mark largely remain, but RAD21 ChIP-seq signal is strongly diminished. (C) Aggregate region analysis of the OCIs in the WAPL/CTCF depletion time course. (D) Hi-C matrix plot for the example in (A) for the triple degron cell line. Bottom row show the differential Hi-C map for the Hi-C matrix plots shown in the two top rows. (E) Aggregate region analysis of the OCIs in the triple degron line.

We hypothesize that in the absence of WAPL and CTCF cohesin can extrude beyond the limits of the OCI. Moving in opposite directions at roughly equal speed will result on average in the observed plume pattern. To test whether plumes are dependent on the (extrusion activity of the) cohesin complex we analyzed the Hi-C data we generated for our triple degron line. Depletion of WAPL/CTCF recapitulates the plume phenotype (**Figure 5D**). Consistent with our hypothesis, depletion of RAD21 following the induction of plumes leads to a rapid loss of plumes (**Figure 5D,E**). These results show that rapid stabilization of cohesin in the absence of CTCF leads to the formation of transient cohesin-dependent plumes. Plumes can span inactive chromatin regions, indicating that in the absence of CTCF cohesin-mediated loop extrusion can invade inactive regions.

### Rapid reorganization in compartmentalization following cohesin stabilization

Within chromosome territories, active and inactive chromatin segregates into distinct microenvironments, called nuclear compartments, which can be identified using Hi-C as megabase-sized compartmental domains (Lieberman-Aiden et al., 2009; Stevens et al., 2017). Knock-out of the cohesin loading factor *Nipbl* or *MAU2* results in increased compartmentalization (Haarhuis et al., 2017; Schwarzer et al., 2017), while stabilization of cohesin by knocking out *WAPL* weakens compartmentalization (Haarhuis et al., 2017; Wutz et al., 2017). However, in these studies compartmentalization was measured after long-term depletion of these cohesin regulators. Using our degron lines we can assess the direct consequences of architectural protein loss on compartmentalization. Following WAPL depletion we find that compartmentalization is already decreased after 6h, although we only see a full diminishment after 24h, with no further decrease at 96h (**Figure 6A and S6A**). Compartmentalization is defined by higher than expected homotypic interactions (i.e. A/A and B/B) and lower than expected heterotypic interactions (i.e. A/B). To quantify the degree of hetero- and homotypic compartment interactions we have performed a saddle plot analysis (**Figure 6B)**. The saddle plots show that WAPL depletion mostly affects the A/A compartment interactions, whereas B/B compartment interactions are less affected. When we deplete WAPL and CTCF simultaneously, compartmentalization is also decreased (**Figure 6A and S6A**). However, when we quantify the changes in interactions between the compartments in the WAPL/CTCF-depleted cells we find that B/B compartment interactions are mostly affected at 6h and 24h, while changes in A/A compartment interactions are much more limited (**Figure 6B**). These results show that changes in compartmentalization can be different depending on the presence of CTCF.

**Figure 6:**
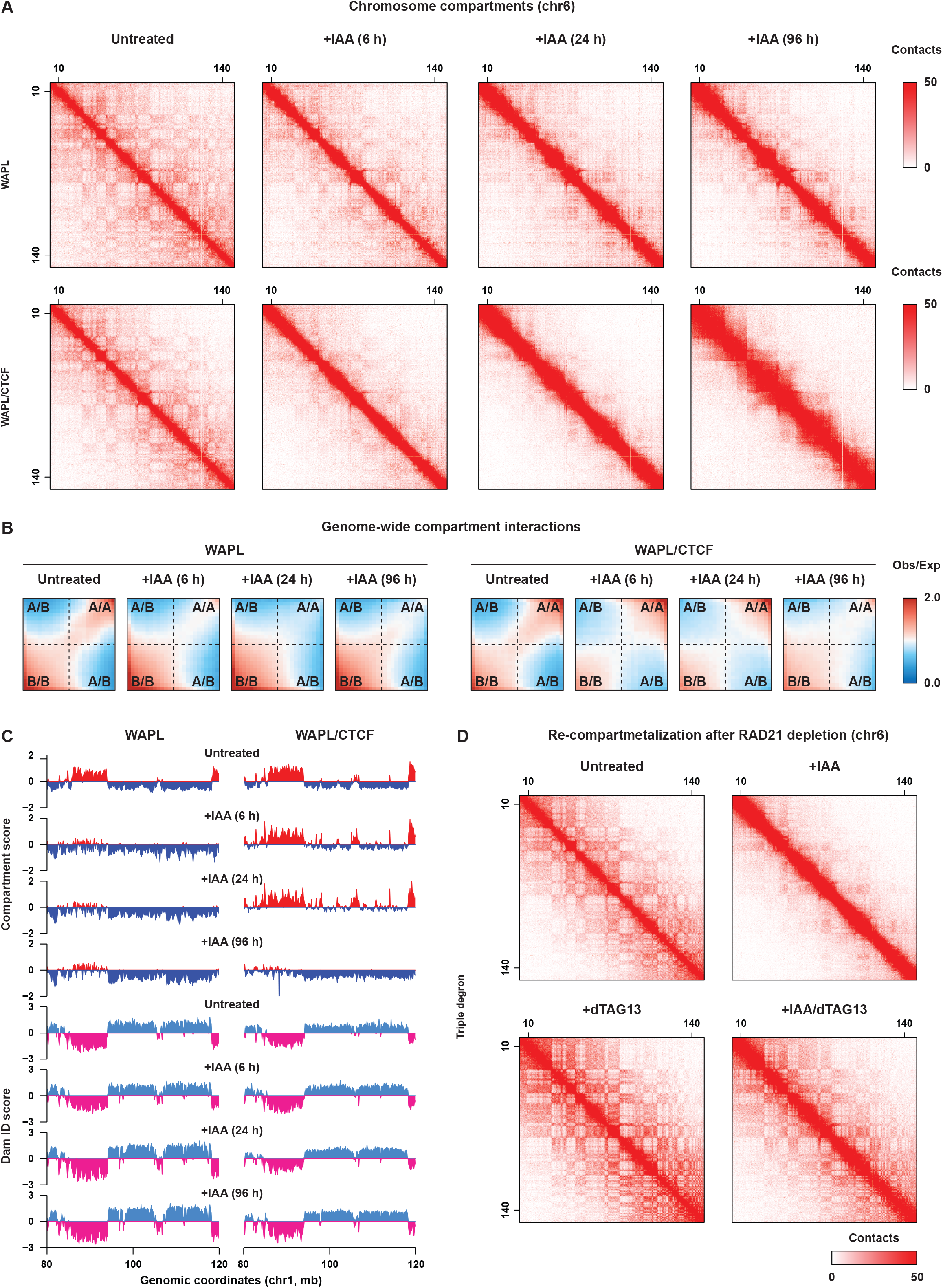
Rapid changes in compartmentalization are driven by cohesin. (A) An example Hi-C matrix plot showing whole chromosome compartment organization at 100kb resolution. (B) Genome-wide saddle plot show strength of compartmentalization. (C) Compartment scores are plotted for a region on chromosome 1 (top four rows). Positive and negative scores indicate A and B compartments, respectively. Bottom four rows show pA-DamID results for Lamin B1 to measure interactions with the nuclear lamina. (D) Chromosome-wide Hi-C matrix plot for the triple degron lines for the treatments indicated.

It has previously been observed that B compartments strongly correlate with lamina associated domains (LADs) and the compartment score shows a strong anti-correlation to the Lamin B1 DamID signal (Kind et al., 2015). Our WAPL and WAPL/CTCF degron systems, which enable us to rapidly induce changes in compartmentalization, are an ideal system to investigate the relationship between compartmentalization and association with the nuclear periphery. However, regular DamID, that is used to measure lamina interactions, requires expression of the Dam-fusion construct for ∼24h (Guelen et al., 2008), precluding the temporal analysis we did for the Hi-C. To solve this we made use of a nuclear lamina interaction dataset generated for a companion study (van Schaik et al., manuscript in preparation) using the recently developed pA-DamID method (van Schaik et al., 2020), which uses antibody-guided *in vitro* Dam methylation and enables taking a snapshot of the genome-lamina interactions. In the untreated cells there is a strong anti-correlation between the compartment score and the LaminB1 signal similar to previous observations (**Figure 6C**). When we compared the compartment scores and LaminB1 DamID signal upon WAPL and WAPL/CTCF depletion, we found that despite the massive changes in compartmentalization, the lamina association seems not to be affected to the same degree (**Figure 6C, Figure S6B**). These results show that nuclear compartmentalization can be uncoupled from nuclear lamina-association, indicating that they are not inherently driven by the same molecular mechanism.

To confirm that the decrease in compartmentalization following the depletion of WAPL is indeed driven by the cohesin complex, we analyzed the compartmentalization of the genome in our triple degron line. Consistent with the proposed role for loop extrusion in preventing compartmentalization, we find that loss of RAD21 increases the compartment strength (**Figure S6C**). Next, we analyzed the compartmentalization in cells in which we first induced a decrease in compartmentalization by depleting WAPL/CTCF followed by RAD21 depletion. We find that in these cells compartmentalization is restored to a wild-type level as indicated by visual inspection of the heatmaps, compartment strength analysis and saddle plot analysis (**Figure 6D, S6C and S6D**). These results show that the cohesin compex is indeed responsible for the restriction of compartmentalization, likely through loop extrusion. Furthermore, it also shows that restoration of physiological levels of compartmentalization occurs on the order of hours, indicating that changes in compartmentalization can be achieved within the span of a single cell cycle.

## DISCUSSION

In this study we assessed to the role of key architectural proteins in 3D genome organization using acute depletion strategies. Acute loss of WAPL recapitulates what we and others have previously described for knock-out or knock-down (Haarhuis et al., 2017; Wutz et al., 2017). By using acute protein degradation techniques that enable the synchronous depletion of a protein, we can now quantitatively evaluate 3D genome reorganization upon loss of architectural proteins with a high temporal resolution. Our observation that loops are reorganized within 6 hours following WAPL depletion confirm earlier studies that the 3D genome is a highly dynamic (Gibcus et al., 2018; Rao et al., 2017). These studies also emphasize the importance of using degron lines to assess the role of architectural proteins in chromosome organization, particularly if one wants to discern direct from indirect effects.

### Acute depletion of cohesin-associated proteins reveals extrusion-dependent structures

The various degron lines presented here enable us to refine the role that cohesin-mediated loop extrusion plays in the establishment of the architectural features of chromosomes. Cohesin-mediated loop extrusion explains why CTCF-anchored chromatin loops are formed between distally located CTCF sites preferentially oriented in a convergent manner (Rao et al., 2014; Vietri Rudan et al., 2015). However, Hi-C data seems to suggest that not all CTCF sites contribute to loop formation. By integrating our cohesin ChIP-seq data with our Hi-C data we now find that regions where cohesin accumulates following WAPL depletion are also the site of the strongest loop anchors. Generally, these are regions with a high density of CTCF sites. Although cohesin is stabilized on chromatin, CTCF likely is not. Dissociation of CTCF would lead to re-initiation of extrusion by cohesin. We propose that when an extruding cohesin complex encounters another cohesin complex they will block each other, explaining why cohesin accumulates at loop anchors and the observation of counter-stripes. This is consistent with the cohesin traffic jam model proposed for loop formation in the absence of WAPL (Allahyar et al., 2018). We expect that in the presence of WAPL, a cohesin complex extruding into a cohesin complex stabilized at a CTCF site will also be stalled, however, the continuous activity of WAPL will resolve these structures before they can be measured.

When we deplete WAPL and CTCF together, extrusion by the cohesin complex is no longer restrained. The plumes, that we observe in this condition may also be explained as a consequence of unrestrained loop extrusion. Consistent with this, we find a strong local enrichment for cohesin peaks within the region of origin of the plume. Acute depletion of WAPL will lead to the synchronized stabilization of cohesin at plume origins. However, in the presence of CTCF, cohesin will remain “trapped” at plume origins. Co-depletion of WAPL and CTCF will lead to the extrusion of cohesin beyond the plume origin and to the juxtaposition of the genomic regions flanking plume origins. This juxtaposition manifests itself as plumes in the Hi-C map. In regions with many open chromatin sites and cohesin binding sites that are flanked by genomic regions that are depleted of by them, cohesin can extrude largely uninterrupted. Intriguingly, plumes are reminiscent of observations made in bacteria. In *Bacillus subtilis*, SMC complexes, which are related to cohesin, can be coordinately loaded at the origin of replication using a temperature sensitive mutant (Banigan et al., 2020; Wang et al., 2017, 2018). Following SMC complex loading, a cross-like organization is revealed by Hi-C. This is explained by a concerted extrusion reaction away from the origin of replication and the “zipping” together of the left and right chromosome arms into a juxtaposed position (Wang et al., 2017).

Depletion of RAD21 following induction of plumes confirms that cohesin and likely extrusion is required for the formation of plumes. Interestingly, a similar genome structure, named “hinge-like” domains or flares, has been observed in zebrafish sperm (Wike et al., 2021). Further analysis is necessary to determine whether plumes and flares are formed by similar or distinct mechanisms.

We and others have previously shown that knock-out of WAPL leads to a decrease in compartmentalization (Haarhuis et al., 2017; Wutz et al., 2017). Our degron lines enabled us to add a temporal component to this loss in compartmentalization. Within 6 hours there is already a severe decrease in compartmentalization. Strikingly, the decrease in inter-compartment interaction is stronger for A compartments than for B compartments when WAPL is depleted. However, when both WAPL and CTCF are depleted this difference is not observed anymore. To explain this difference, we need to consider that the vast majority of cohesin complexes are found in A compartments. Upon WAPL depletion these complexes accumulate at CTCF binding sites, of which the majority is also found in A compartments. Therefore, cohesin complexes will be largely restricted from extruding in the B compartment. However, when CTCF is depleted together with WAPL, cohesin will no longer accumulate at CTCF binding sites. Our plume analysis has shown that cohesin can likely extrude into flanking regions of low gene activity (i.e. B compartments). We propose that this extrusion into B compartmental domains disrupts the interactions between B compartments (**Figure 7**).

**Figure 7:**
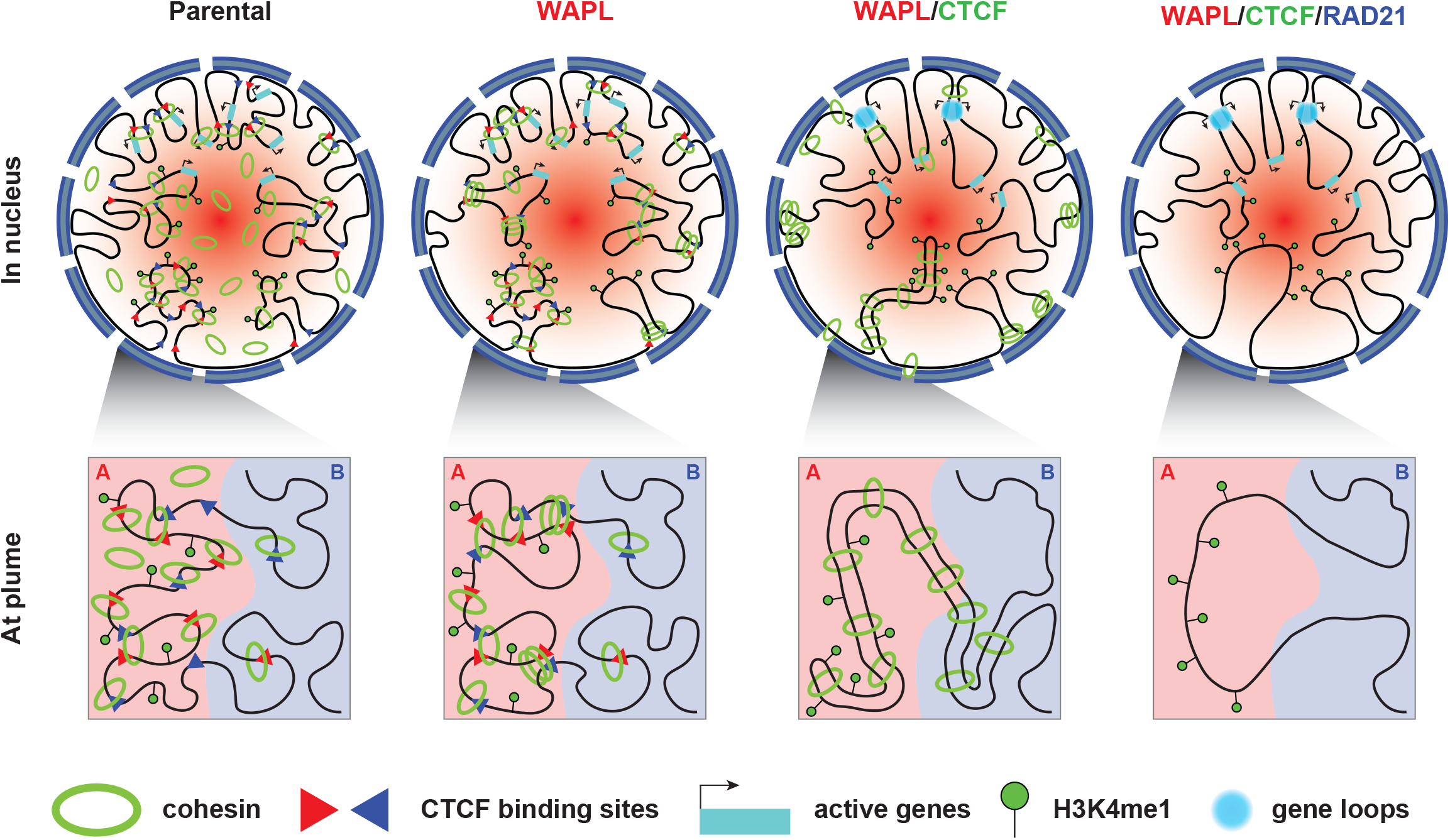
A model summarizing 3D genome organization changes after acute depletion of key architectural factors. The upper panel shows the higher-order genome organization in one nucleus in the parental, WAPL-deficient, WAPL/CTCF-deficient, or WAPL/CTCF/RAD21-deficient cells. The lower panel describes spatial DNA arrangement at an architectural plume in the four cell lines.

We show that disruption of intercompartment interactions has only mild effects on the position of compartmental domains with respect to the nuclear periphery. Given the strong anti-correlation between the LaminB1 DamID score and compartment scores, the formation of compartments and LADs could be driven by similar mechanism. However, WAPL depletion, which drastically affects the compartment score, has only mild effects on lamina association of B compartments. Although surprising, this in not wholly unexpected. In the 2-cell mouse embryonic stage there is an absence of compartments, but single cell DamID experiments have shown that there are regions that are specifically associated with the nuclear lamina (i.e. LADs) (Borsos et al., 2019). However, this is first time that we have, in a system where both LADs and compartments are both formed, disrupted one without severely affecting the other. Although we find only mild effects on the large-scale organization of LADs, in a companion paper we have looked at more subtle consequences of changes in cohesin positioning on chromosome interactions with the nuclear lamina (van Schaik et al., manuscript in preparation).

### Interactions between active genes are not mediated by cohesin

When we deplete both WAPL and CTCF new loops start to emerge. We find that these loops are strongly associated with actively transcribed genes. Loss of WAPL and CTCF has been shown to lead to an accumulation of cohesin at sites of convergent transcription, dubbed ‘cohesin islands’ (Busslinger et al., 2017). Although we observed the same phenomenon, the effect is not as strong, likely because mESCs are rapidly cycling, even in the absence of WAPL (Liu et al., 2021) or CTCF (Nora et al., 2017). Breakdown of the cohesin complex during mitosis ensures the reloading and therefore repositioning of the cohesin complex in the next cell cycle. What has remained unclear was whether cohesin positioning at these active genes contributes to nuclear organization. Surprisingly, in our sequential depletion analysis, we found no evidence that active genes or RNA PolII can act as cohesin-mediated loop anchors analogous to CTCF. Under normal WAPL and CTCF expressing conditions cohesin dynamically forms short-range interactions (<1Mb), preventing mis-aggregation of active genes over a large genomic distance. However, when cohesin is stabilized in the absence of CTCF, much longer interactions are formed, promoting the interactions between active genes. In the absence of cohesin these interactions are preserved, indicating that although extrusion is involved in the formation of these loops it is dispensable for their maintenance.

What can be the mechanism that drives the preferential association between active genes? Both Hi-C (Bonev et al., 2017) and ChIA-PET (Li et al., 2012) analyses have revealed clustering of active promoters. Live cell imaging experiments have further revealed that RNA PolII can form droplet-like structures inside the nucleus (Cisse et al., 2013), suggesting that a phase-separation mediated mechanism may underlie the clustering of RNA PolII. Whatever, the mechanism, by looping together distally located active genes, cohesin may facilitate the self-association of RNA PolII. In parallel, other active chromatin features, such as H3K36me3 or associated proteins may be involved.

Our results clearly show the importance of using acute depletion methods for studying chromosome organization, which led us to identify 3D genome configurations that are transient (i.e. architectural plumes) and study proteins whose loss of function induces a cellular state change (e.g. WAPL) (Liu et al., 2021). In addition, we show that by using two degron techniques in a sequential manner we can dissect the molecular mechanisms that drive chromosome architecture. Finally, the increased temporal resolution afforded by acute protein degradation enables us to uncouple highly correlated architectural features to show that these features are unlikely to be formed by the same mechanism. In the pursuit of elucidating the role of the 3D genome in processes such as DNA replication, DNA repair and gene regulation acute protein depletion will play a crucial role.

## STAR★METHODS

### Key Resources Table

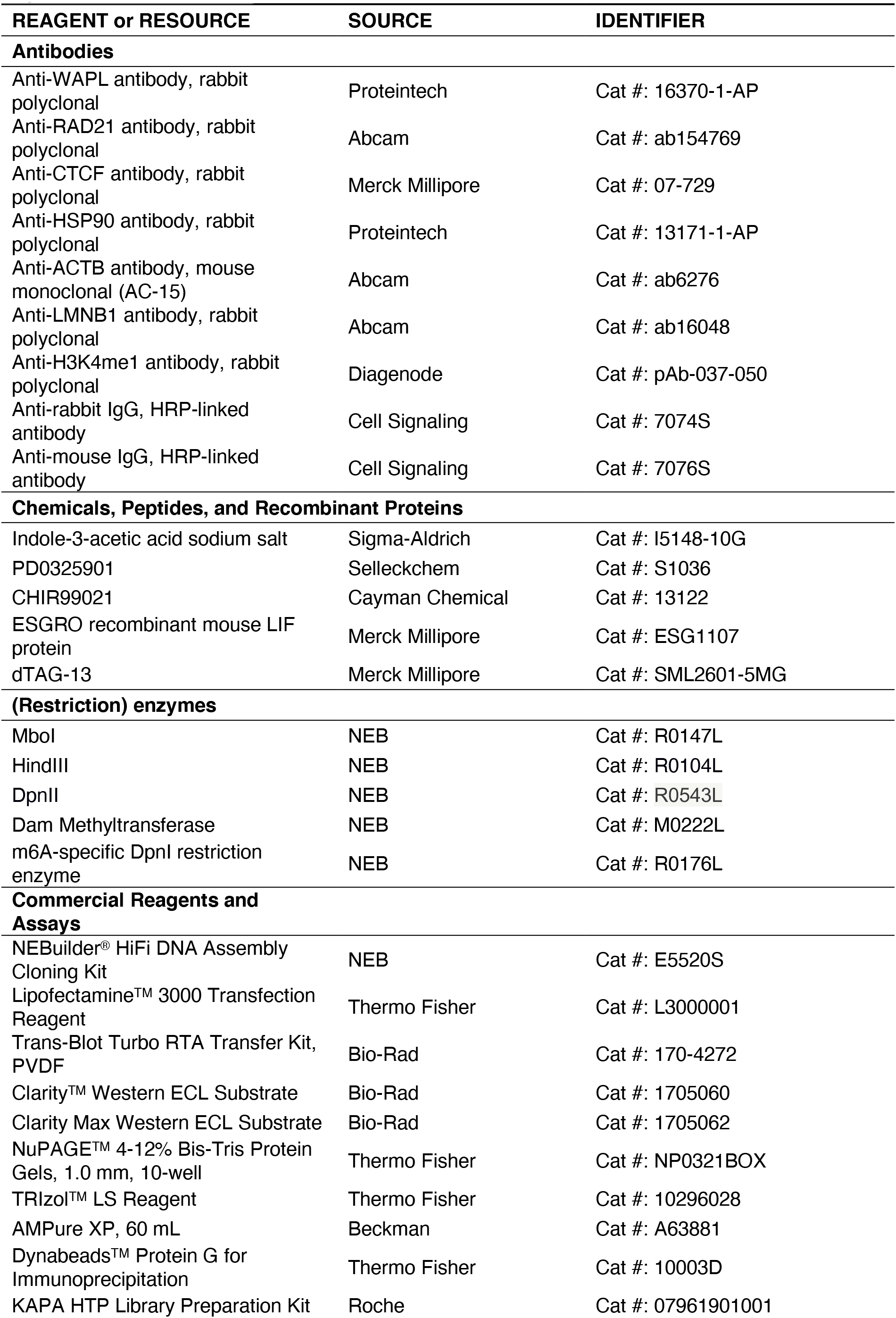
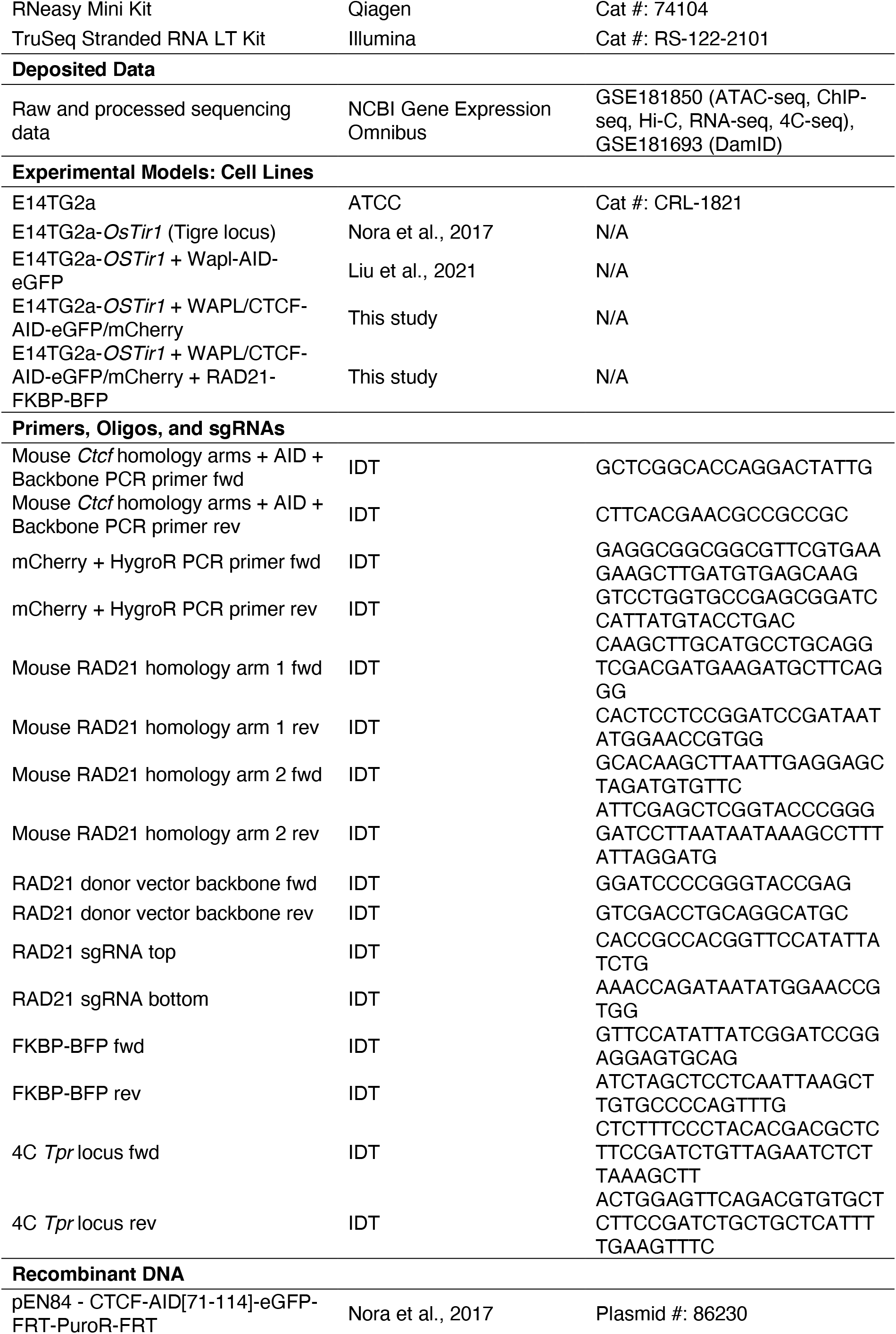
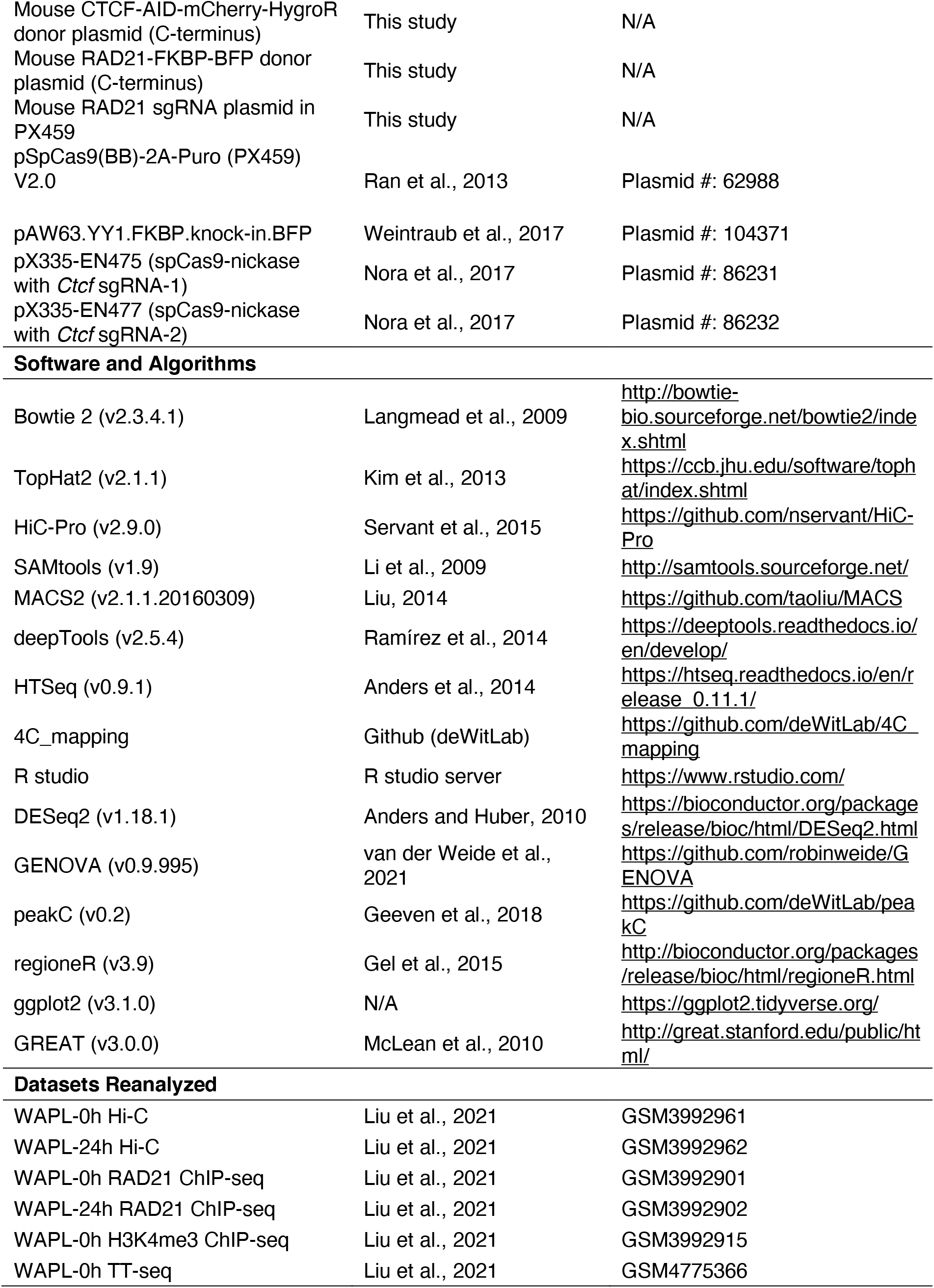

### Contact for Reagent and Resource Sharing

Further information and requests for reagents, plasmids and cell lines should be directed to the Lead Contact, Elzo de Wit.

### Experimental Model and Subject Details

#### Mouse Embryonic Stem Cells (ESCs)

E14Tg2a (129/Ola isogenic background) and the degron tagged cell lines were cultured on 0.1% gelatin-coated plates in serum-free DMEM/F12 (Gibco) and Neurobasal (Gibco) medium (1:1) supplemented with N-2 (Gibco), B-27 (Gibco), BSA (0.05%, Gibco), 10^4^ U of Leukemia Inhibitory Factor/LIF (Millipore), MEK inhibitor PD0325901 (1 μM, Selleckchem), GSK3-β inhibitor CHIR99021 (3 µM, Cayman Chemical) and 1-Thioglycerol (1.5×10^-4^ M, Sigma-Aldrich). The cell lines were passaged every 2 days in daily culture. During the protein depletion experiments, the cells were seeded overnight before the start of the time course in the following densities: For a 96 h time course, 35 k, 150 k, and 400 k cells were seeded in 6-well, 10-cm and 15-cm plates, respectively. For 24 h or 27 h time course, 0.5 M, and 4 M cells were seeded in 6-well and 15-cm plates, respectively. The medium was refreshed or the cells were split in 1:10 every 2 days during a time course.

#### Acute protein degradation and inhibition

The WAPL and CTCF/WAPL proteins were (co-)depleted by treating the cells with a final concentration of 500 μM IAA (Sigma-Aldrich). The RAD21 protein was depleted by adding a final concentration of 500 nM dTAG-13 molecule (Merck Millipore) (Nabet et al., 2018).

All the time series experiments were performed by inducing protein degradation at different time points and harvest the samples in the end of the time course.

### Method Details

#### Plasmid Construction

The donor plasmid used to target the endogenous mouse CTCF and RAD21 protein was constructed by modifying a published pEN84 plasmid (Addgene). To construct the *Ctcf* donor plasmid, two homology arms around the stop codon of the *Ctcf* genes, a backbone and an AID peptide were co-amplified by PCR from the pEN84 plasmid. A mCherry fluorescent protein sequence and a hydromycin resistant protein sequence were co-amplified from an in-house available plasmid (Mouse WAPL-AID-mCherry-HygroR donor plasmid). The amplified DNA fragments were joint into a mouse CTCF-AID-mCherry-HygroR donor plasmid using the NEBuilder^®^ HiFi DNA Assembly Cloning Kit (NEB). To construct the *Rad21* donor plasmid, the HA-P2A sequence was first removed from the pAW63 (Addgene). Subsequently, we amplified two homology arms around the C-terminus of the *RAD21* gene and the FKBP-BFP sequence and the backbone from the modified pAW63 plasmid. The PCR products were then assembled using the NEBuilder^®^ HiFi DNA Assembly Cloning Kit (NEB).

The donor sequences and sgRNAs in the obtained plasmids were validated by Sanger sequencing before using for further experiments.

### Gene Targeting

Generation of the CTCF/WAPL-AID cell line was performed in the background of our WAPL-AID cell line (Liu et al., 2021). The donor plasmids and their corresponding sgRNAs (pX335-EN475 and pX335-EN477, Addgene) for *Ctcf* targeting were co-transfected into the parental cell lines using Lipofectamine 3000 Reagent (Thermo Fisher). Three days after transfection, the eGFP/mCherry double positive cells were sorted into a gelatinized 96-well plate for single clone selection. The obtained clones were genotyped by PCR and the fusion sequences were validated by Sanger sequencing. The CTCF/WAPL-AID+RAD21-FKBP cell line was further generated in the background of the CTCF/WAPL-AID cell line. For *Rad21* targeting, the donor plasmid and the sgRNA (pX330-EN1082, Addgene) were electroporated into the CTCF/WAPL cells using Neon Transfection System (Thermo Fisher). We sorted eGFP/mCherry/BFP triple positive cells and perform single clone selection and validation following the same strategy as described above.

### Western Blots

We extracted nuclear soluble fraction of the mESCs for blotting the WAPL, CTCF, POLR2A and ACTB proteins. For RAD21 and HSP90, mESCs were harvested and lysed in RIPA lysis buffer (150 mM NaCl, 1% NP-40, 0.5% sodium deoxycholate, 0.1% SDS, and 25 mM Tris (pH=7.4)).

The NuPAGE^TM^ 4-12% Bis-Tris Protein Gels (Thermo Fisher) were used to separate the proteins. The separated protein was transferred to a pre-activated PVDF membrane using Trans-Blot Turbo Transfer System (Bio-Rad). The blots were incubated with the following primary antibodies overnight at 4°C: (1) WAPL (1:1000, 16370-1-AP, Proteintech), (2) RAD21 (1:1000, ab154769, Abcam), (3) CTCF (1:1000, 07-729, Merck Millipore), (4) POLR2A (1:2000, 39097, Active Motif), (5) ACTB (1:5000, AC-15, Abcam) and (6) HSP90 (1:2000, 13171-1-AP). After incubation, the blots were washed 3 times with TBS-0.1% Tween-20. The blots were then incubated with secondary antibody against rabbit IgG at room temperature for 1 h, following by 3-time TBS-0.1% Tween-20 washing. The proteins attached with antibodies were hybridized with Clarity^TM^ Western ECL Substrate or Clarity Max Western ECL Substrate reagent (Bio-Rad) and visualized in a ChemiDoc MP Imaging System (Bio-Rad).

### ChIP-seq

The ChIP-seq experiments were performed in presence of 10% HEK293T cells as an internal reference using a published protocol with small modifications (Liu et al., 2017). For chromatin preparation, the mouse embryonic stem cells were mixed with 10% HEK293T cells and cross-linked by a final concentration of 1% formaldehyde for 10 min. The cross-linking reaction was quenched using 2.0 M glycine. The cross-linked cells were then lysed and sonicated to obtain ∼300 bp chromatin using Bioruptor Plus sonication device (Diagenode). For ChIP assays, antibodies were first coupled with Protein G beads (Thermo Fisher), and then the sonicated chromatin were incubated overnight at 4°C with the antibody coupled Protein G beads. After over incubation, captured chromatin was washed, eluted and de-crosslinked. The released DNA fragments were purified using MiniElute PCR Purification Kit (Qiagen). The ChIP experiments were performed using the following antibodies: (1) WAPL (16370-1-AP, Proteintech), (2) CTCF (07-729, Merck Millipore), (3) RAD21 (ab154769, Abcam), and (4) H3K4me1 (pAb-037-050, Diagenode).

The purified DNA fragments were prepared according to the protocol of KAPA HTP Library Preparation Kit (Roche) prior to sequencing. All the ChIP-seq libraries were sequenced using the single-end 65-cycle mode on an Illumina HiSeq 2500.

### RNA-seq

RNA was isolated following a standard TRIzol RNA isolation protocol. The cells were lysed using 1 ml of TRIzol^TM^ LS Reagent (Thermo Fisher), and 200 μL chloroform was added to the lysates. The mixture was vortexed and centrifuged at 12,000 g at 4°C for 15 min. Upper phase was homogenized with 0.5 ml of 100% isopropanol, incubated at room temperature for 10 min, and centrifuged at 4°C for 10 min. The resulted RNA pellet was washed with 75% ice-cold ethanol, dried at room temperature for 10 min, and resuspended in RNase-free water. The isolated RNA was treated with DNase using RNeasy Mini Kit (Qiagen).

RNA-seq libraries were prepared using a TruSeq Stranded RNA LT Kit (Illumina). The libraries were sequenced using the same platform as the ChIP-seq libraries.

### Hi-C

We generated Hi-C data as previously described (Rao et al., 2014) with minor modifications (Haarhuis et al., 2017). For each template, 10 million cells were harvested and crosslinked using 2% formaldehyde. Crosslinked DNA was digested in nucleus using MboI (NEB), and biotinylated nucleotides were incorporated at the restriction overhangs and joined by blunt-end ligation. The ligated DNA was enriched in a streptavidin pull-down. Hi-C libraries were prepared using a standard end-repair and A-tailing method and sequenced on an Illumina HiSeq X sequencer generating paired-end 150 bp reads.

### 4C-seq

We generated 4C data for untreated and 24h IAA treated Wapl-AID and Rad21-AID cells. 4C was performed as previously described (Geeven et al., 2018; van de Werken et al., 2012) using a two-step PCR method for indexing described first in (Haarhuis et al., 2017). We used HindIII (NEB) as the first restriction enzyme and DpnII (NEB) as the second restriction fragment. Viewpoint specific primers can be found in the section of STAR&Methods. The 4C-seq libraries were sequenced using the same platform as the ChIP-seq libraries.

### pA-DamID

pA-DamID experiment was performed using our published protocol (van Schaik et al., 2020). Briefly, two million cells of each experimental condition were harvested and kept on ice. One million cell was used to localize pA-Dam to a Lamin B1 antibody (1:100, ab16048, Abcam). The remaining cells were used as Dam-control by adding 0.5 μL free Dam Methyltransferase (NEB) during incubation with methyl donor S-adenosylmethionine. Genomic DNA was digested with the m6A-specific DpnI restriction enzyme (NEB) and further processed for high-throughput sequencing. Dam-control and Lamin B1 samples were sequenced on an Illumina HiSeq 2500 with approximately 30 million 65 bps single-end reads per sample.

### ATAC-seq

ATAC-seq experiment was performed following a published protocol (Liu et al., 2017). We first isolated the nuclei from the harvested mESCs, and then the nuclei were permeabilized and tagmented using in-house-generated Tn5 transposase. The tagmented DNA was amplified by two sequential nine-cycle PCR amplificaitons, and the DNA fragments between 150 bp and 700 bp in size were purified with AMPure XP SPRI beads (Beckman). The ATAC-seq library was sequenced on a NextSeq 550 platform.

### Quantification and Statistical Analysis

#### ChIP-seq Analysis

Calibrated ChIP-seq data were analyzed using our published pipeline (Liu et al., 2021). Raw sequencing data was mapped to a concatenated reference genome (mm10 and hg19) using Bowtie 2 mapper (version 2.3.4.1) (Langmead et al., 2009). The spike-in reference (reads mapped to hg19) was used for the calibration in order to properly quantify the samples (reads mapped to mm10).

### ChIP-seq Peak Alignment and Functional Annotation

Alignment of ChIP-seq signal was performed using deepTools v3.0 (Ramírez et al., 2016). The “Scale-regions” method was applied to align the signal coverage from broad regions. The “reference-point” method was used for alignment to slim regions (TSSs and insulation borders). Heatmaps were directly made using deepTools. Alignment plots were generally made with aligned matrices that were further processed in R.

### CTCF Motif Analysis

Peaks were identified based on the CTCF ChIP-seq data. Again, peaks with at least 10 reads were kept for further analyses. Fimo v4.11.2 was used to locate CTCF motifs (MA0139.1 from the JASPER database) in these CTCF ChIP-seq peaks (Grant, 2011).

### RNA-seq Analysis

Raw RNA-seq data were mapped against mm10 reference genome using a TopHat2 pipeline (version 2.1.1) (Kim et al., 2013). The mapped reads with mapping quality score <10 were discarded using SAMtools. The read coverage for each gene in “Mus Musculus GRCm38.92” annotation file was determined using a HTSeq tool (version 0.9.1). The coverage files were generated using “normalize to 1X genome coverage” methods in deepTools.

### ATAC-seq analysis

ATAC-seq data were were mapped against mm10 reference genome using BWA-MEM (version 0.7.15-r1140)(Li and Wren, 2014). The mapped reads with mapping quality score <15, as well as optical PCR duplicates, were discarded using SAMtools. The coverage files were generated using “normalize to 1X genome coverage” methods in deepTools (version 3.0).

### Hi-C data Processing

Raw Hi-C data were mapped with HiC-Pro (version 2.9.0) (Servant et al., 2015), which performs mapping, identification of valid Hi-C pairs, generation of contact matrices and ICE normalization (Imakaev et al., 2012). Subsequent analyses were performed in GENOVA, a Hi-C visualization tool written in R (http://github.com/deWitLab/GENOVA) (Van Der Weide et al., 2021). We used the primary loops as identified in earlier studies, with more deeply sequenced data (Bonev, 2017). Based on the loop anchors of these primary loops, we identified putative extended loops by recombining 5’ and 3’ loop anchors, with distance limits of 1.5 to 3 Mb.

To split the Aggregate Peak Analyses for overlap between the loop anchors and RSCs we used the GenomicRanges package. To determine the overlap of RSCs with loop anchors, we reduced the loop anchors to single set of non-overlapping regions. This set was used to determine whether an individual RSC overlaps with a loop anchor. Similar to the extended loops, pair-wise interactions were created between RSCs. For each RSC, we identified overlaps with CTCF motifs (see Motif Analysis section). We selected only those RCSs that overlap with motifs in a single orientation. We then generated pair-wise interactions with only these RSCs.

### 4C-seq Analysis

The raw sequence data was mapped using our 4C mapping pipeline (http://github.com/deWitLab/4C_mapping). We normalized our 4C data to 1 million intrachromosomal reads and visualize chromatin interactions around the viewpoints using peakC (http://github.com/deWitLab/peakC) (Geeven et al., 2018).

### Loop calling

We used Chromosight v1.4.1 (Matthey-Doret et al., 2020) for *de novo* loop annotation for WAPL-AID and WAPL/CTCF-AID cell lines. Loops were identified for 20 kb resolution Hi-C maps for untreated and merged protein depleted time points. For loop detection the Pearson correlation threshold was set to 0.45 and 0.4 for WAPL-AID and WAPL/CTCF-AID respectively, loop sizes were set between 100 kb and 10 Mb, and parameter --smooth-trend was enabled. Loop anchors were classified as shared, if within ±1 bin there was a loop anchor from another condition, and condition-specific otherwise. Permutation test with circular randomization from regioneR (Gel et al., 2016) was used to calculate the enrichment of overlaps of ChromHMM states for mESC (Pintacuda et al., 2017) with the loop anchors. Background enrichment was assessed using shifted loops obtained by randomly shifting *de novo* annotated loops between 100 kb and 1 Mb.

Aggregate Peak Analyses of the original and *de novo* loops were performed using z-score transformed Hi-C matrices at 20 kb resolution.

### Plume annotations

To annotate the plumes, we used the peaks based on ATAC-seq in the untreated condition (see ATAC-seq analysis). For every 100 kb bin, we summed up the scores of the peaks it overlaps with. This gives us a score per bin. If the combined score of the bin and its 4 flanking bins is over 100, we consider the bin to be in a region with high ATAC scores. We set a limit to the maximum size of an ATAC island at 10 consecutive bins of high ATAC scores, i.e. a maximum stretch of 1Mb. Additionally, we required the flanking 1Mb on both sides to be devoid of bins in a high ATAC region. To assess RAD21 ChIP-seq signal at the plumes and their flanking region, we calculated the ChIP-seq peaks per MB for the plumes and the flanking 2Mb bins on either side of the plume.

### Compartment Analysis

We calculated the compartment score using GENOVA (van der Weide, 2021). The orientation for the compartment score for the untreated condition was determined using H3K4me1 ChIP-seq data. The later timepoints are oriented in such a way to give the highest correlation with the compartment score from the untreated sample. Saddle plots are created with GENOVA. Briefly, the saddle plot is used to illustrate the abundance of inter- vs intra-compartment interactions. Interactions between regions of based on the compartment score quantiles are used to do so.

### pA-DamID Analysis

The DamID adapter was trimmed from the 65 bp single-end reads using Cutadapt (version 1.11) and custom scripts. The remaining gDNA was mapped to mm10 with BWA-MEM (version 0.7.17). Further processing was done with custom R scripts. Reads overlapping GATC fragment ends (indicating successful DpnI digestion) were counted into 100kb bins and subsequently normalized. These bins were first normalized to 1M reads and with a pseudocount of 1 a log2-ratio over the Dam-only control was calculated. We used the 100kb resolution for the downstream comparison with the compartment scores. At least two biological replicates were performed for every experimental condition. The average signal between replicates was used for downstream analyses.

## ACKNOWLEDGMENTS

We thank the NKI Genomics Core Facility for help with sequencing and the NKI Flow Cytometry Facility for help with single-cell sorting of genome-edited cells. We thank Teun van den Brand for valuable suggestions for data analysis. We thank Dr. Elphège P. Nora for providing the OsTIR1 parental cell line for our genomic editing. We thank Dr. Boyan Bonev for sharing the coordinates of chromatin loops detected from the high-resolution Hi-C data generated from the wild-type E14Tg2a cell line for our analyses. We thank members of the de Wit lab for critically reading the manuscript. Work in the de Wit laboratory is supported by an ERC StG 637587 (‘HAP-PHEN’), an ERC CoG 865459 (‘FuncDis3D’) and a Vidi grant from the Netherlands Scientific Organization (NWO, 016.16.316). N.Q.L. is supported by a Veni grant from the Netherlands Scientific Organization (NWO, 016.Veni.181.014). The de Wit and van Steensel laboratories are part of Oncode, which is partly financed by the Dutch Cancer Society.

## CONTRIBUTIONS

N.Q.L. and E.d.W. conceived and designed the study. N.Q.L., and H.T. performed experiments in the laboratories of E.d.W., and T.v.S. performed the DamID analysis under the supervision of B.v.S.. N.Q.L., M.M., M.M.G.A.S., T.v.S., R.H.v.d.W., and E.d.W. analyzed data. E.d.W. supervised the study. N.Q.L. and E.d.W. wrote the manuscript with input from all authors.

---

**Figure S1:**
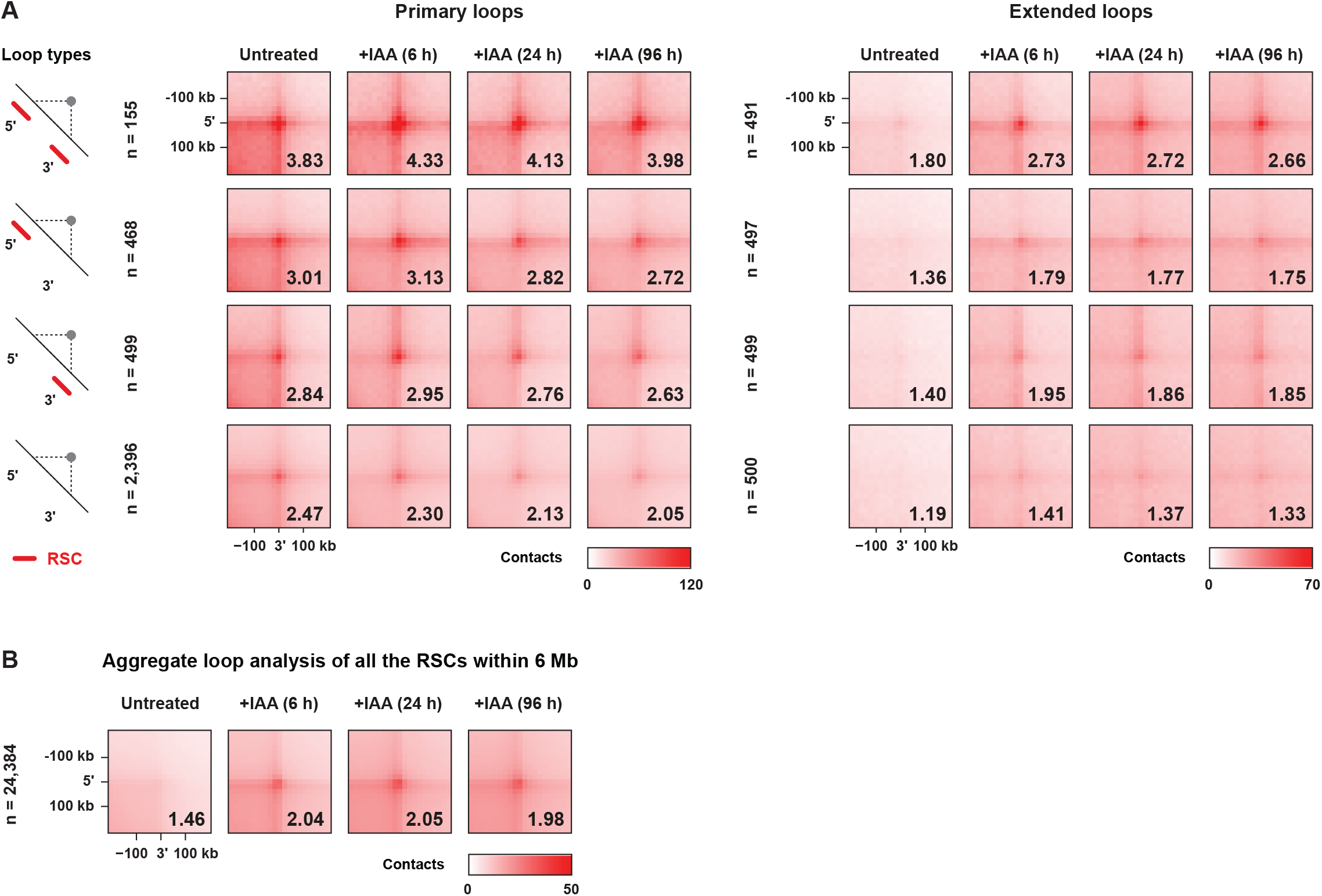
Aggregate peak analysis of loop characteristics following WAPL depletion. (A) Primary and extended loops are quantified using aggregate peak analysis. Types of the loops are stratified by the presence of an RSC at the loop anchor. (B) Genome-wide quantification of putative, *in silico* generated, chromatin loops formed by two RSCs within 6Mb (randomly sampled loops from the entire list).

**Figure S2:**
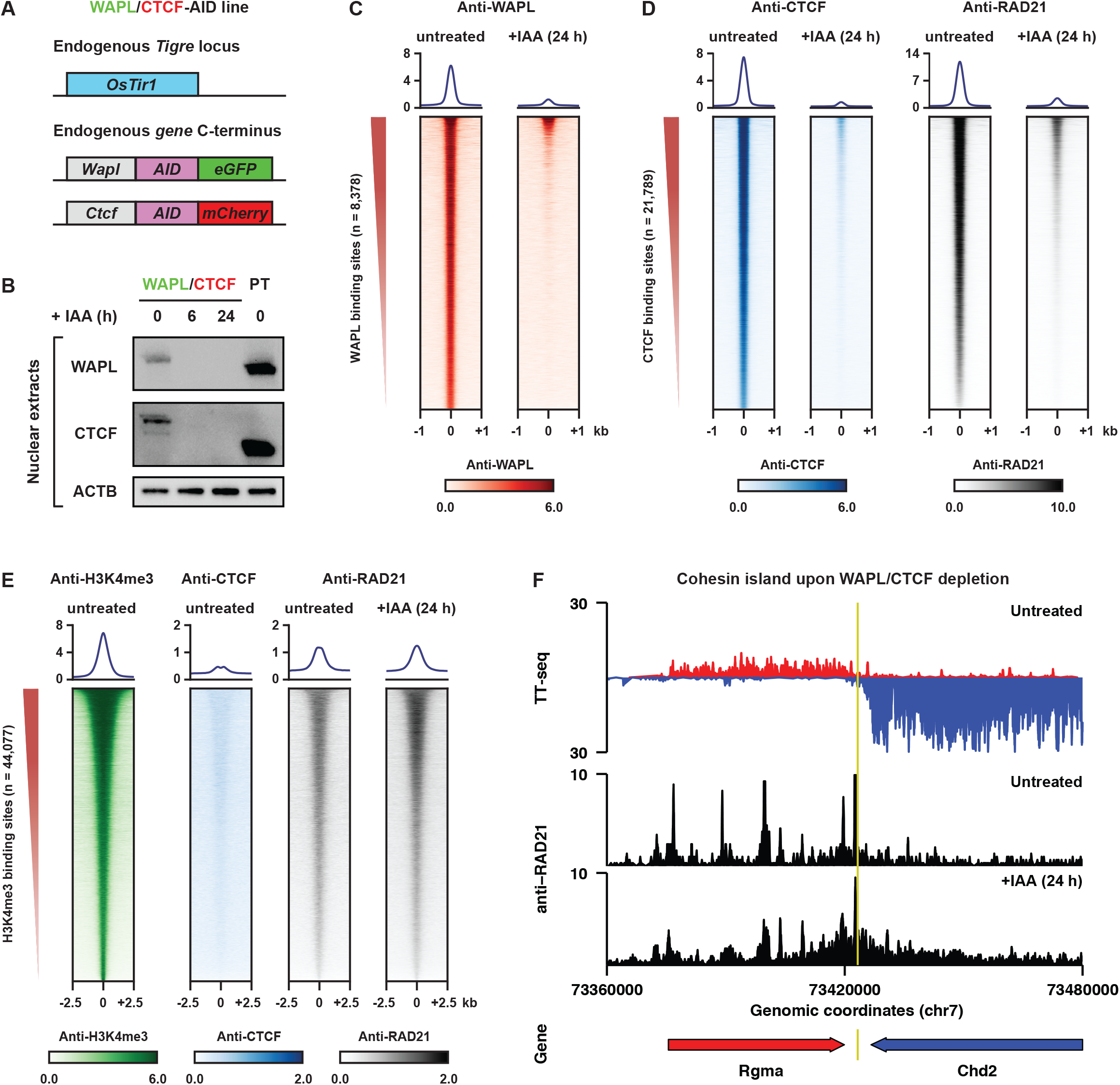
Molecular characterization of the WAPL/CTCF-AID cell line. (A) In the OsTir1 E14 mESCs, the endogenous *Wapl* and *Ctcf* gene were edited to create AID–eGFP and AID-mCherry fusions at the C-terminus of WAPL and CTCF, respectively. (B) Western blot analysis validates acute depletion of WAPL and CTCF upon IAA treatment. Tornado plots summarize calibrated ChIP-seq analyses for WAPL (C), CTCF and RAD21 (D) before and after IAA treatment. (E) Tornado plot quantifying RAD21 binding at H3K4me3 binding sites (active TSSs). (F) Results of TTseq in untreated WAPL-AID cells and RAD21 ChIPseq in treated and untreated WAPL/CTCF-AID cells at a “cohesin island” described by Busslinger et al. (2017).

**Figure S3:**
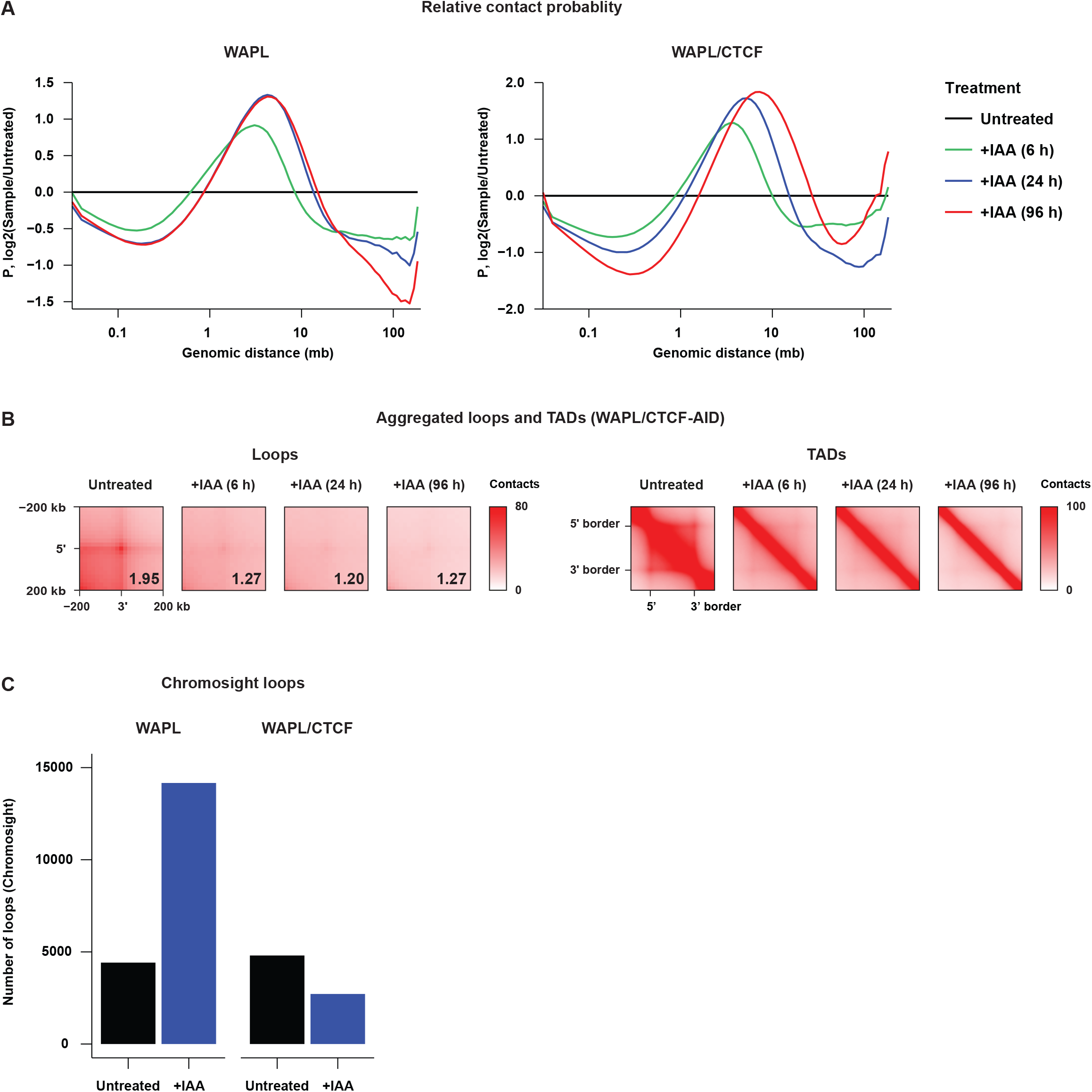
Genome-wide changes in the 3D genome following WAPL/CTCF depletion. (A) Relative contact probability following WAPL and WAPL/CTCF depletion shown relative to the untreated samples. (B) Aggregate TAD and loop analysis for WAPL/CTCF depletion time course. Loops and TADs are the same as shown in Figure 1B and are obtained from Bonev et al. (2017). (C) Number of chromatin loops detected by ChromoSight loop caller in the WAPL and WAPL/CTCF depletion experiments.

**Figure S4:**
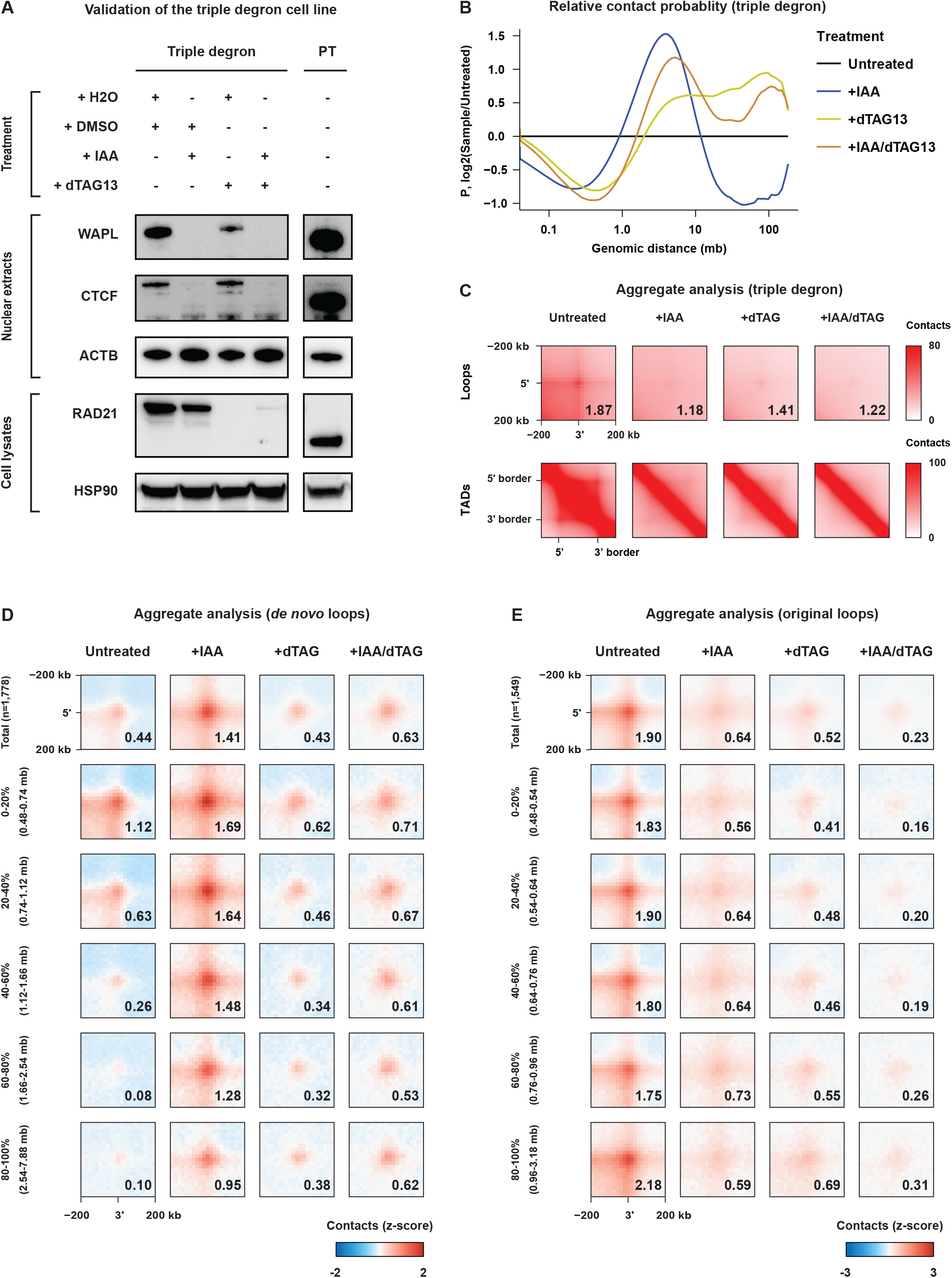
Generation of the triple degron cell line and genome-wide effects on 3D genome organization. (A) Western blot analysis confirms sequential depletion of WAPL/CTCF and RAD21 after IAA and dTAG13 treatment, respectively/ (B) Relative contact probabilities in the triple depletion experiment following different treatments; results shown relative to untreated cells. (C) Aggregate TAD and loop analysis for different triple degron treatment conditions. The loops and TADs are the same as shown in Figure 1B and are obtained from Bonev et al. (2017). (D) Aggregate loop analysis on z-score normalized Hi-C matrices of the triple degron lines with different treatments for WAPL/CTCF *de novo* loops. Loops are stratified into five equally size bins and aggregate signal is calculated per size bin. (E) Same as (D) but for loops identified in the WAPL/CTCF untreated Hi-C data (original loops).

**Figure S5:**
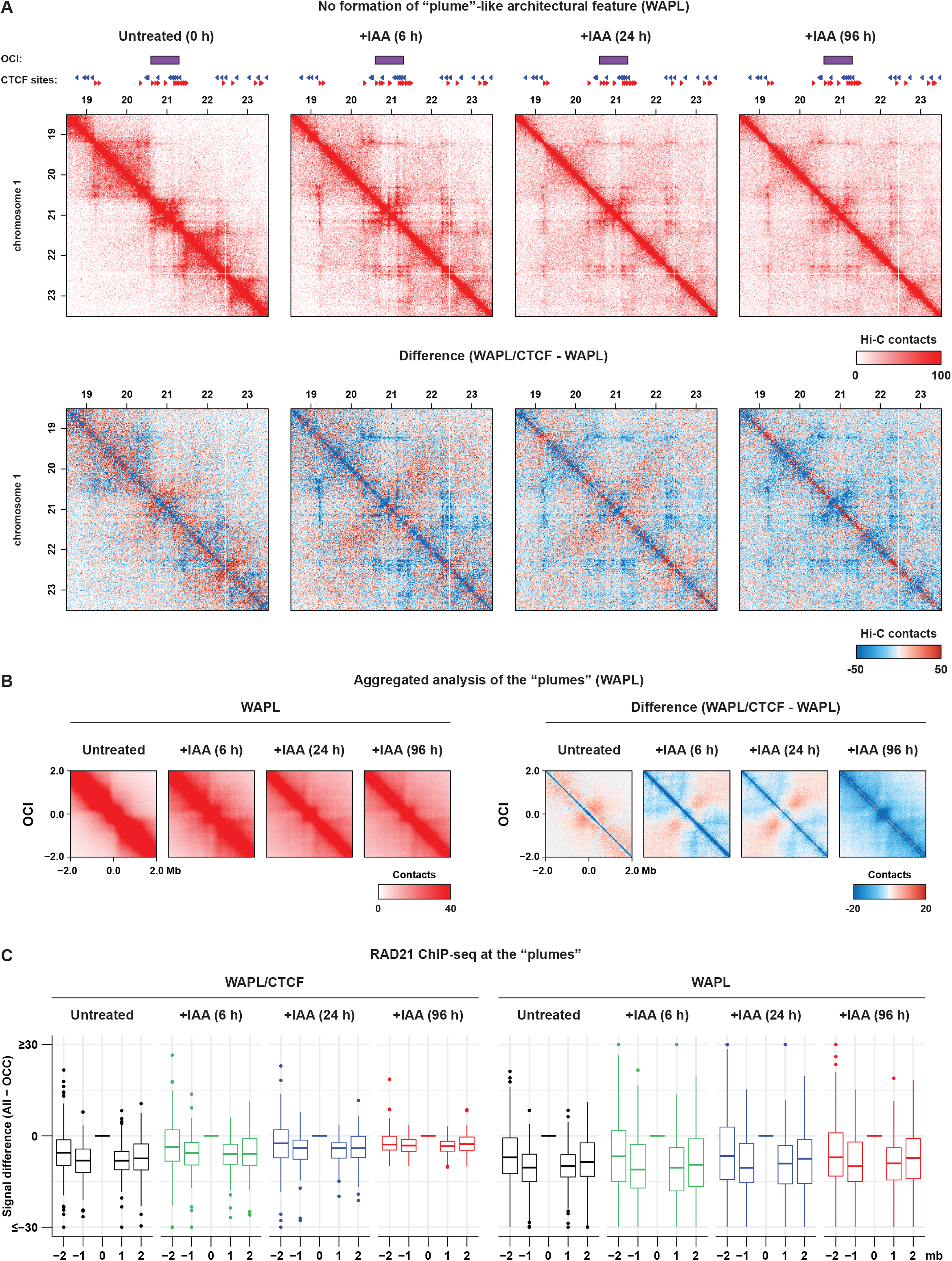
WAPL/CTCF-double depletion is required to form the “plume”-like architectural features. (A) Top row shows Hi-C matrix plot at 20kb resolution for the WAPL-AID time course showing the example region from Figure 5A. Bottom row show the difference between WAPL/CTCF depletion Hi-C data and WAPL only depletion Hi-C data. (B) Aggregate region analysis (ARA) using the OCIs for the WAPL-AID time course Hi-C data (left panel). Right panel shows difference between WAPL/CTCF and WAPL only depletion ARA data for the OCIs in the left panel. (C) Boxplot showing the average RAD21 ChIPseq signal in the OCI (middle box) and the flanking regions. Average RAD21 signal is defined as average calibrated ChIPseq coverage calculated using deepTools.

**Figure S6:**
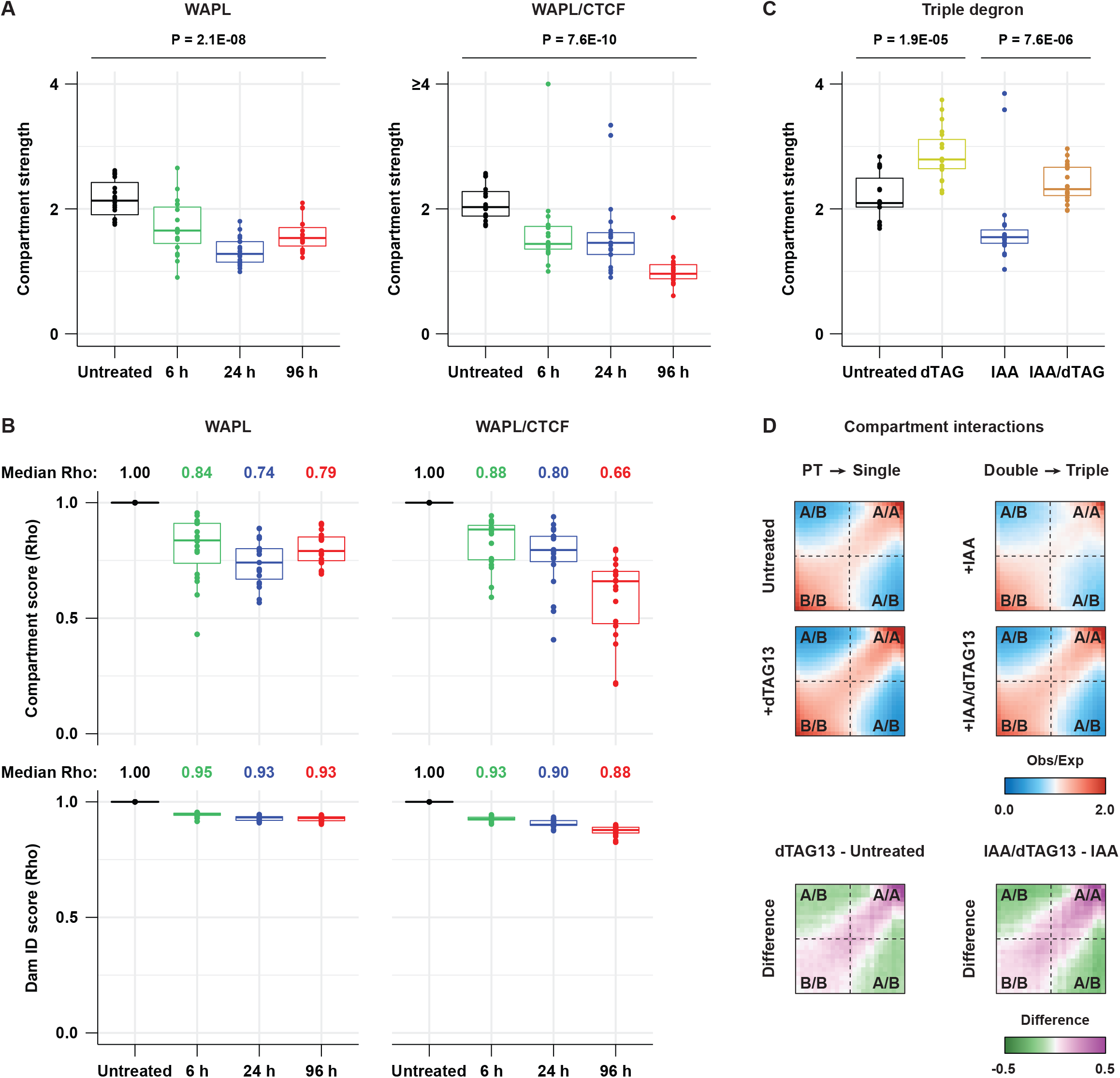
Genome-wide quantification of the changes in compartment strength and association with nuclear lamina interactions. (A) Genome-wide quantification of compartment strength, defined as the ratio of the homotypic (A/A or B/B) over heterotypic (A/B) interactions. Individual dots show the compartment strength per chromosome. (B) Spearman correlation coefficients showing the (dis)similarity of compartment scores for untreated Hi-C data or Hi-C data from time points following treatment (top panels). Bottom panels shows same as top panel but for Spearman correlation coefficients of LaminB1 DamID data. Individual dots show individual chromosomes. (C) Genome-wide quantification of the compartment strength in the triple degron line following different treatments. (D) Genome-wide saddle analysis for the Hi-C data from the triple degron line following different treatments. Bottom panels show differential saddle plots following dTAG treatment in either untreated or WAPL/CTCF depleted cells.

